# Cohesin Facilitates Nucleosome Invasion by Transcription Factors

**DOI:** 10.64898/2025.12.08.693039

**Authors:** Shane Stoeber, Hengye Chen, Ruo-Wen Chen, Courtney Smith, Sam Becker, Sarah Vinson, Michael G. Poirier, Lu Bai

## Abstract

Nucleosomes present a major barrier to transcription factor (TF) binding. However, a subgroup of TFs known as pioneer factors (PFs) can recognize motifs covered by nucleosomes and initiate chromatin opening. PFs also bind nucleosomal substrates with high affinity *in vitro*, which may facilitate their nucleosome invasion *in vivo*. Here, we show that LexA, a bacterial TF with poor nucleosome binding *in vitro*, can rapidly invade into a well-positioned nucleosome from a motif positioned at the dyad when expressed ectopically in yeast. This striking contrast between LexA binding *in vitro* and *in vivo* raises the possibility that TFs can exploit nucleosome dynamics *in vivo* to access occluded sites. Surprisingly, we find that LexA-mediated chromatin opening can occur in the absence of DNA replication, chromatin remodeling, histone turnover, and a few histone chaperones. Instead, nucleosome invasion by LexA and a native PF, Cbf1, is promoted by the cohesin complex, illustrating intriguing a connection between cohesin and TF binding. Together, our results demonstrate that even non-pioneer TFs like LexA can bind and displace nucleosomes *in vivo*, through a process facilitated by cohesin.

## INTRODUCTION

Transcription factors (TFs) play a central role in orchestrating gene expression programs. Understanding the factors that promote or inhibit TF binding is therefore critical in elucidating the mechanisms of gene regulation. TFs typically function by recognizing specific DNA sequence motifs, which are often embedded within nucleosomes in eukaryotic genomes. Because histone proteins make extensive contacts with DNA, nucleosome-embedded motifs are less accessible, significantly restricting the ability of most TFs to engage with their target sites and regulate transcription^1–3^. Over the past few decades, a specialized class of TFs called pioneer factors (PFs) has emerged. These PFs possess the unique ability to engage with nucleosomal substrates, increase chromatin accessibility, and facilitate the binding of downstream TFs that would otherwise be excluded by the nucleosome barrier^4–6^. In higher eukaryotes, PFs are often regarded as “master regulators” for differentiation and cell-fate commitment. For example, OCT4 and SOX2 are essential in maintaining embryonic stem cell identity, and FOXA1/2 are required for proper endoderm and hepatic development and differentiation^7,8^.

Despite extensive research, identifying the biochemical properties of TFs that confer pioneering activity remains a topic of debate^9,10^. Many biochemical studies of purified PFs, such as Reb1, Cbf1 in yeast, FOXA1, OCT4, SOX2 in mammals, and Zelda in *Drosophila*, have shown that these factors exhibit similar binding affinities for naked and nucleosomal DNA^4–6,11,12^. Some PFs achieve this using a “dissociation rate compensation” mechanism, in which they dissociate more slowly from nucleosomal DNA through interactions with histones to enhance their binding affinity towards nucleosomal substrates^11,13,14^. Inside cells, PFs display slow nuclear mobility and the ability to scan and access closed chromatin^15^. Together, these studies support a model where PFs possess special intrinsic biochemical properties that allow them to effectively engage nucleosomes, which eventually leads to nucleosome displacement and/or histone eviction.

In contrast, other evidence suggests that such intrinsic nucleosome-binding properties are not strictly necessary for pioneering activity. Nucleosome displacement results from TFs and histones competing for the same genomic locus, and this competition can be modulated by the TF concentration. For example, a yeast TF Pho4 binds nucleosomes poorly and lacks dissociation rate compensation *in vitro*, yet it can displace nucleosomes when overexpressed in yeast^13^. Rfx1, another yeast TF with weak pioneering activity, can be converted into a strong PF at elevated expression levels^16^. In mammalian cells, HNF4A is typically classified as a non-PF because it binds downstream of FOXA1 during hepatocyte differentiation and preferentially targets open, DNase-hypersensitive sites^17,18^. However, when overexpressed in human cells, HNF4A can bind and open previously inaccessible chromatin regions^19,20^. These observations indicate that sequence-specific TFs with high concentration may be sufficient to confer pioneering activity, raising the possibility that any TF can function as a PF in certain conditions.

One caveat of these *in vivo* experiments is that TFs are assessed in their endogenous cellular context, where their pioneering activities can potentially be affected by interactions with other TFs and/or co-regulators. Moreover, studies in mammalian cells typically rely on ATAC-seq to infer chromatin accessibility, a method that lacks single-nucleosome resolution. Even when nucleosome-level resolution is achieved with Micrococcal Nuclease (MNase) mapping, most nucleosomes in mammalian genomes adopt fuzzy positioning^21^, making it difficult to evaluate the exact location of TF motifs within the nucleosome.

To address these limitations, we employed a minimal system by expressing the *E. coli* TF LexA in budding yeast to investigate its nucleosome-displacing activity. LexA is a small transcriptional repressor (∼23 kDa) that binds as a dimer to a palindromic consensus motif (LexO) with high affinity (*K*_d_ ∼1 nM)^3^. It is composed of an N-terminal helix-turn-helix (HTH) DNA binding domain along with a C-terminal dimerization domain. Structural studies of the LexA-DNA complex reveal that LexA binding induces DNA bending towards LexA, a configuration that is incompatible with nucleosomes^22^. Consistent with this idea, LexA was shown to bind poorly to nucleosomes^23^. When the LexO is placed at 8 bp from the entry side of a Widom 601 nucleosome, LexA binding rate is reduced by two orders of magnitude, while its dissociation rate is increased up to three orders of magnitude, resulting in dramatically lowered binding affinity compared to naked DNA^3^. As a bacterial TF, it is unlikely for LexA to have specific coordinated interactions with euchromatic chromatin regulators. Therefore, using this artificial system, we can ask whether a TF with strong DNA binding but weak nucleosome binding can access its nucleosome-embedded motif *in vivo*, and if yes, which cellular processes promote such activity.

Here, we show that LexA can efficiently and rapidly invade into nucleosomes and generate a local nucleosome-depleted region (NDR) in yeast, even when its motif is positioned at the dyad of a well-positioned nucleosome. As this process likely relies on nucleosome dynamics *in vivo*, we systematically measured LexA-mediated nucleosome invasion in the absence of factors typically associated with nucleosome dynamics. Surprisingly, we find that chromatin opening by LexA occurs independently of DNA replication, chromatin remodeling, and a few histone chaperones. Instead, this process is inhibited by histone turnover and facilitated by the cohesin complex. Cohesin also promotes nucleosome invasion and chromatin binding by endogenous yeast PF Cbf1. These findings reveal an unanticipated connection between cohesin, nucleosome dynamics, chromatin accessibility, and TF binding.

## RESULTS

### LexA binds weakly to nucleosomes *in vitro* but invades into nucleosomes rapidly *in vivo*

Previous studies of LexA-nucleosome interactions used the Widom 601 nucleosome positioning element^3^. Nucleosomes assembled on 601 are highly stable and may pose an unusually strong barrier for LexA binding. To determine whether LexA’s nucleosome binding property also applies to naturally-occurring nucleosomes, we examined LexA binding to a yeast native nucleosome, YNN1, which is significantly less stable than 601^24^. Based on previously determined nucleosome positioning over YNN1^24^, we placed LexO at the dyad (position 64-83 bp) and measured LexA binding using electrophoretic mobility shift assay (EMSA). Consistent with previous reports^3^, LexA shows strong binding to LexO on naked DNA with a dissociation constant *K*_D_ < 5 nM (**Figure 1A & B**). However, when the same DNA is reconstituted into nucleosomes, no detectable band shift occurs until LexA concentration reaches 300 nM (**Figure 1A & B**). The same EMSA pattern occurs on nucleosomes lacking LexO sites, indicating that the band shift is due to nonspecific binding. These results confirm that LexA exhibits poor nucleosome binding *in vitro*, even when using a biologically relevant, less stable nucleosome substrate.

**Figure 1.**
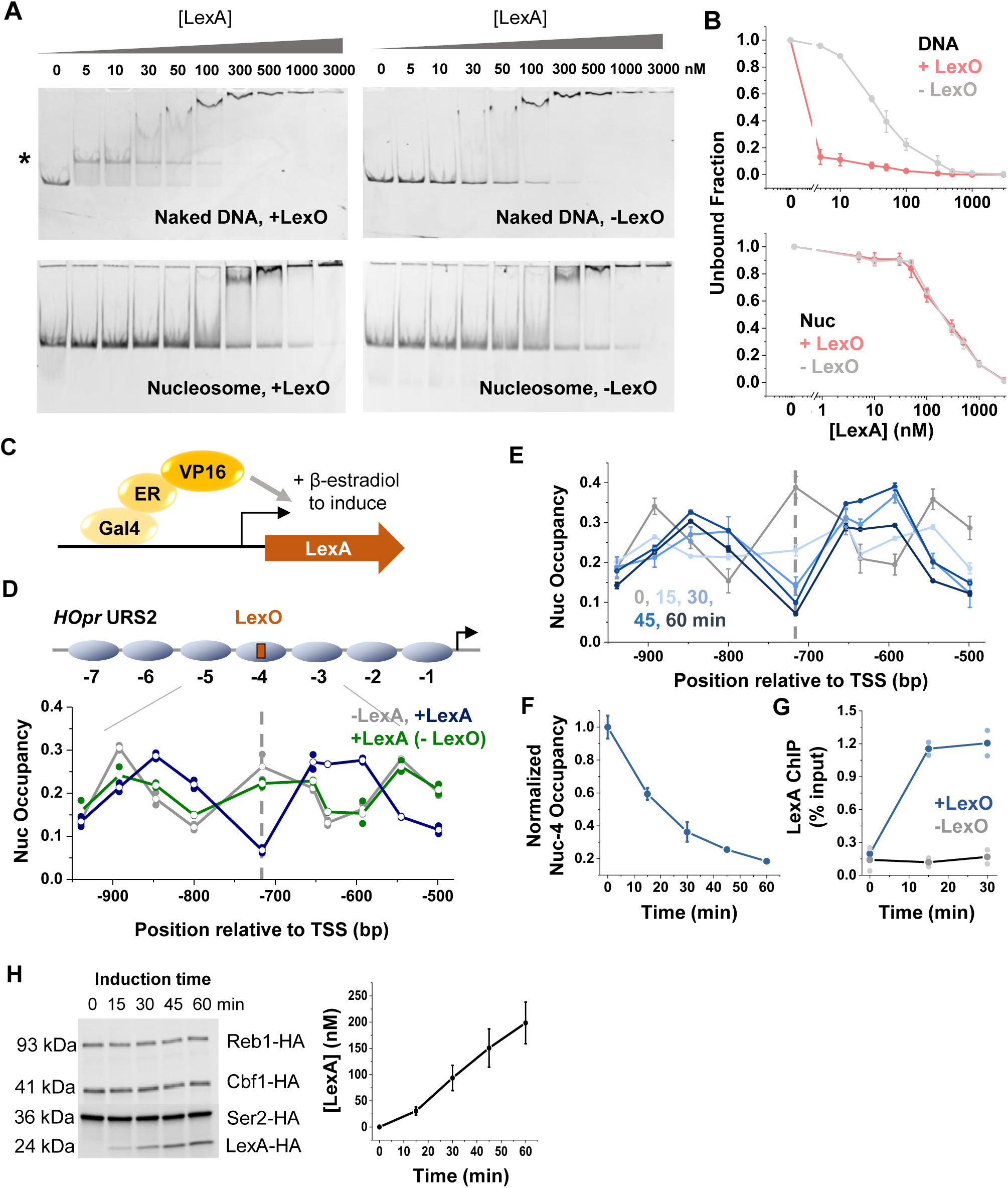
LexA binds poorly to nucleosomes *in vitro* but invades rapidly into nucleosomes in vivo. **A)** Electrophoretic mobility shift assay (EMSA) of LexA on naked or nucleosomal YNN1 DNA containing a LexO motif at the nucleosome dyad (left) or no LexO motif (right). 0.5 nM naked DNA (top) or 0.5nM nucleosomes (bottom) were incubated with increasing amounts of purified LexA. The asterisk indicates the band corresponding to LexA bound to free DNA. **B)** Quantification of LexA EMSA in A. Intensities of unbound DNA bands (top) and unbound nucleosome bands (bottom) are shown. Error bars represent the standard errors from three biological replicates (same for below unless otherwise noted). The presence of LexO does not alter the gel shift pattern on nucleosomes, indicating the lack of motif-specific binding. **C)** Schematic of the construct used to induce LexA expression. **D)** LexA invades into nucleosomes *in vivo* through LexO motif at the dyad of nucleosome –4. Top: nucleosome configurations of a modified *HOpr* with the engineered LexO. Bottom: nucleosome occupancy measured by MNase-qPCR before (gray) and after (blue) LexA induction, as well as over a background *HOpr* sequence lacking LexO after LexA induction (green). The dashed line marks the LexO location (same as below). Two biological replicates were analyzed for each condition. **E)** Time course of LexA invasion into *HOpr* nucleosome –4. MNase-qPCR were performed at different time points during LexA induction. Darker colors represent longer induction times. Three biological replicates were analyzed per time point. **F)** Quantification of the change in nucleosome occupancy at the LexO site from the time course in E. **G)** LexA ChIP-qPCR at nucleosome –4 during LexA induction, over LexO-containing (blue) and LexO-lacking (gray) templates. Two biological replicates were analyzed per time point. **H)** Quantification of LexA induction level. Anti-HA Western blot was performed using a strain expressing HA-tagged Reb1, Cbf1, Ser2 and inducible LexA-HA. LexA-HA levels were estimated by comparing its band intensities with those of the HA-tagged endogenous proteins (see Methods). Two biological replicates were used for each time point.

Next, we assessed LexA binding and nucleosome invasion *in vivo* by inducing LexA expression with 17β-estradiol in a *S.cerevisiae* strain (**Figure 1C**)^25^. This strain also harbors a single LexO site engineered into a well-positioned nucleosome –4 within a modified *HO* promoter (*HOpr*) (**Figure 1D**). The Swi5 binding sites in this *HOpr* are mutated to maintain a constitutive nucleosome array across the cell cycle^16,26^. In the absence of LexA, MNase mapping confirms that LexO resides at the dyad of nucleosome –4 (**Figure 1D & S1A**). After LexA induction, nucleosome occupancy near LexO undergoes a drastic change: nucleosome –4 dyad is converted into a short linker, flanked by two nucleosomes that become completely out of phase with the original array (**Figure 1D**). This effect is dependent on LexO, as LexA induction does not alter nucleosome positioning on the *HOpr* background (**Figure 1D**). These results indicate that LexA can invade into a nucleosome via a single motif at the dyad and displace adjacent nucleosomes *in vivo*.

To measure the dynamics of LexA invasion, we carried out MNase measurements over *HOpr* following LexA induction at different time points(**Figure 1E**). Nucleosome –4 occupancy is altered within 15 min of LexA induction, and its depletion is nearly complete by 30 min (**Figure 1F**). We further monitored LexA binding over time by chromatin immunoprecipitation (ChIP), which detected significant LexA binding at 15 min of LexA induction (**Figure 1G**), consistent with the rapid depletion of nucleosome –4. To test if such nucleosome invasion activity is a result of exceptionally high LexA concentration in cells, we used Western blotting to quantify the concentration of HA-tagged LexA throughout induction. By comparing the band intensities of LexA-HA to three HA-tagged endogenous proteins of known concentrations, we estimated LexA level to be ∼30 nM at 15 min and ∼100 nM at 30 min (**Materials and Methods; Figure 1H, S1B & C**). Notably, LexA does not show nucleosome binding *in vitro* at these concentrations (**Figure 1A**).

Given this apparent contradiction, we asked if the rotational setting of the LexO in **Figure 1D** happens to promote LexA invasion into nucleosome –4. To test this, we repositioned LexO by shifting it 3 bp from its original location (p64 to p67), which will significantly alter its rotational orientation on the nucleosome –4. Similar nucleosome displacement was observed regardless of the LexO positioning (**Figure S1D & E**), indicating that the rotational setting has minimal impact on LexA invasion. Collectively, these findings demonstrate that LexA binds poorly to nucleosomes *in vitro* but can efficiently invade nucleosomes *in vivo*.

### Depletion of chromatin remodelers does not affect the rate of nucleosome invasion by LexA

We hypothesized that the discrepancy between LexA’s behavior *in vitro* and *in vivo* may be due to differences in nucleosome dynamics. Under physiologically relevant salt conditions, purified nucleosomes are relatively stable, exhibiting limited unwrapping or repositioning^27,28^. In contrast, nucleosomes in living cells are subject to constant perturbation by various regulatory factors. In particular, chromatin remodelers (CRs) use energy derived from ATP-hydrolysis to translocate nucleosomes or evict histones from DNA^29^. This raises the possibility that CR-dependent nucleosome dynamics may transiently expose otherwise occluded motifs and facilitate TF binding (**Figure 2A**).

**Figure 2.**
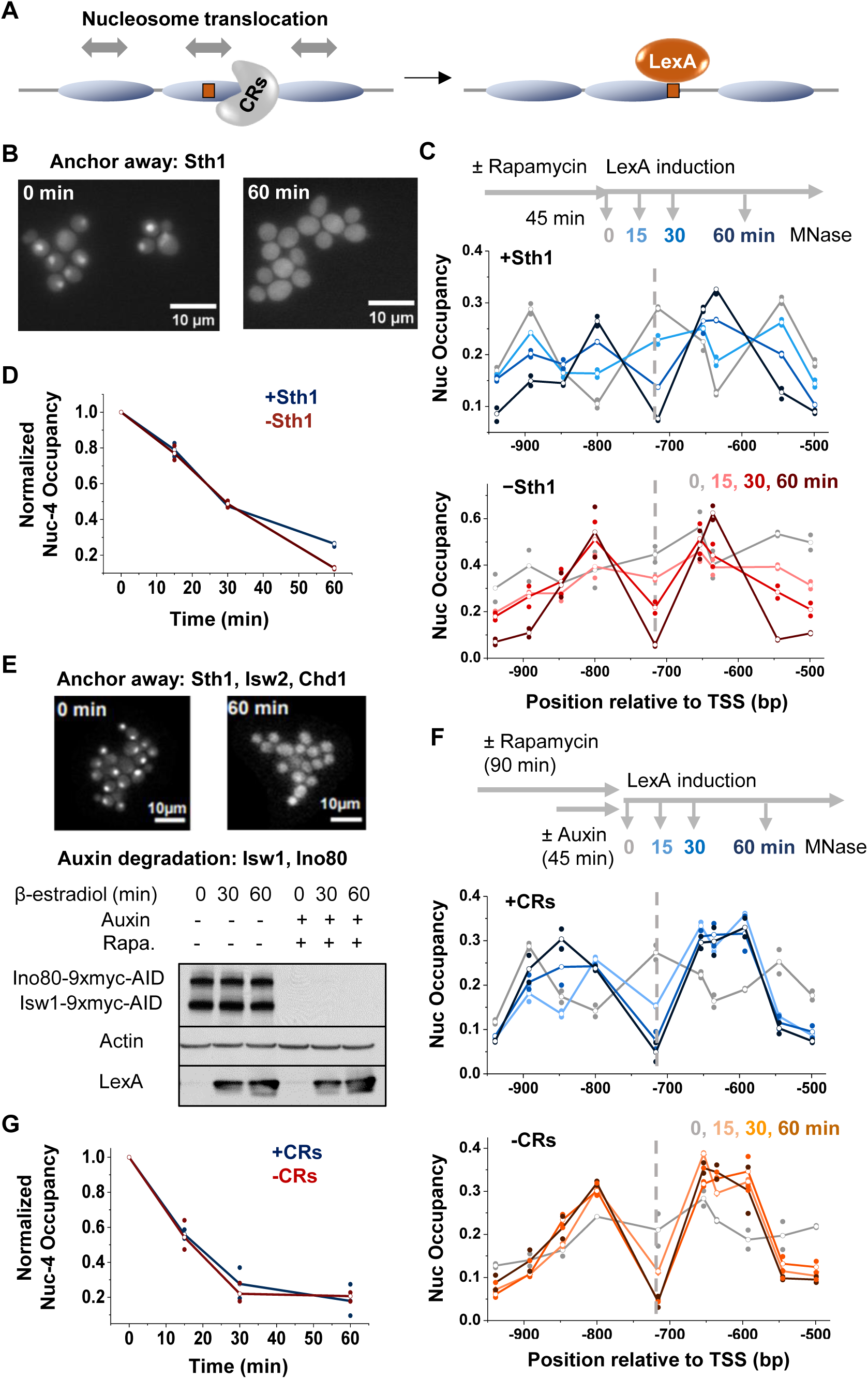
The rate of LexA invasion is not affected by depletion of chromatin remodelers (CRs). **A)** Cartoon illustrating how the ATP-dependent remodeling activity of CRs could facilitate nucleosome displacement and promote LexA binding. **B)** Sth1 anchor away. Fluorescent images of yeast cells expressing Sth1-GFP before (left) and after (right) rapamycin treatment confirm nuclear depletion of Sth1. **C)** Time course of LexA invasion into nucleosome –4 in the presence (top) or absence of Sth1 (bottom). **D)** Quantification of nucleosome occupancy at the LexO site during LexA induction with or without Sth1. **E)** Simultaneous depletion of multiple CRs. Fluorescence imaging confirms concurrent anchor away of Sth1-GFP, Isw2-GFP, and Chd1-GFP, and a Western blot shows the auxin-induced degradation of Ino80 and Isw1. Actin serves as a loading control. LexA is induced to similar levels regardless of CR depletion. **F)** Time course of LexA invasion into nucleosome –4 in the presence (top) or absence of multiple CRs (bottom). **G)** Quantification of nucleosome occupancy at the LexO site during LexA induction with or without multiple CRs.

To determine whether nucleosome invasion by LexA depends on CR activity, we first examined the role of RSC, the most abundant CR in budding yeast and the only CR essential for cell viability^30^. Depletion of the ATPase subunit of RSC, Sth1, results in genome-wide shrinkage of NDRs, indicating that RSC broadly promotes chromatin accessibility^31–33^ and could, in principle, facilitate LexA invasion. We generated an anchor-away strain to rapidly deplete GFP-tagged Sth1. Fluorescent imaging confirms efficient removal of Sth1 from the nucleus following rapamycin treatment (**Figure 2B**). We then assessed nucleosome occupancy during LexA induction with and without Sth1. Sth1 depletion results in fuzzy nucleosome positioning over the *HOpr* (**Figure 2C**, -Sth1 0 min), consistent with altered remodeling activity. However, LexA-mediated nucleosome displacement remains unchanged, with comparable levels of nucleosome depletion at 15, 30, and 60 min of LexA induction in the presence or absence of Sth1 (**Figure 2C & D**). These findings indicate that RSC is not required for LexA invasion into nucleosomes.

Since different CRs may function redundantly, we constructed another strain to simultaneously remove five major remodelers (Sth1, Isw1, Isw2, Chd1, and Ino80) through a combination of anchor-away and auxin-induced degradation^32^ (**Figure 2E**). Removal of these CRs has no impact on LexA induction levels (**Figure 2E**). Similar to Sth1 depletion alone, the elimination of all five CRs leads to fuzzy nucleosome positioning over the *HOpr* (**Figure 2F**). Despite the loss of most of the remodeling activities in the cell, LexA still efficiently displaces nucleosomes near the LexO site with no detectable changes in the rate of nucleosome invasion (**Figure 2F & G**). Altogether, these results illustrate that LexA-induced nucleosome displacement occurs independently of most ATP-dependent chromatin remodelers.

### Depletion of histone chaperones does not affect the rate of nucleosome invasion by LexA

Another group of proteins that is associated with nucleosome dynamics is histone chaperones, which bind histones and facilitate nucleosome assembly and disassembly independently of ATP hydrolysis^34^. In particular, we focused on FACT, an essential complex known to bind nucleosomes and induce major conformational changes^35^. More specifically, FACT is involved in histone “recycling”, where DNA is temporarily unwrapped from histones and then reassembled^36^ (**Figure 3A**). This process has the potential to expose underlying DNA and contribute to nucleosome displacement. FACT is also highly abundant in yeast, enabling its widespread function in transcription, replication, and DNA repair^37^. Based on this prior knowledge, we reasoned that FACT could be involved in facilitating LexA-mediated nucleosome displacement.

**Figure 3.**
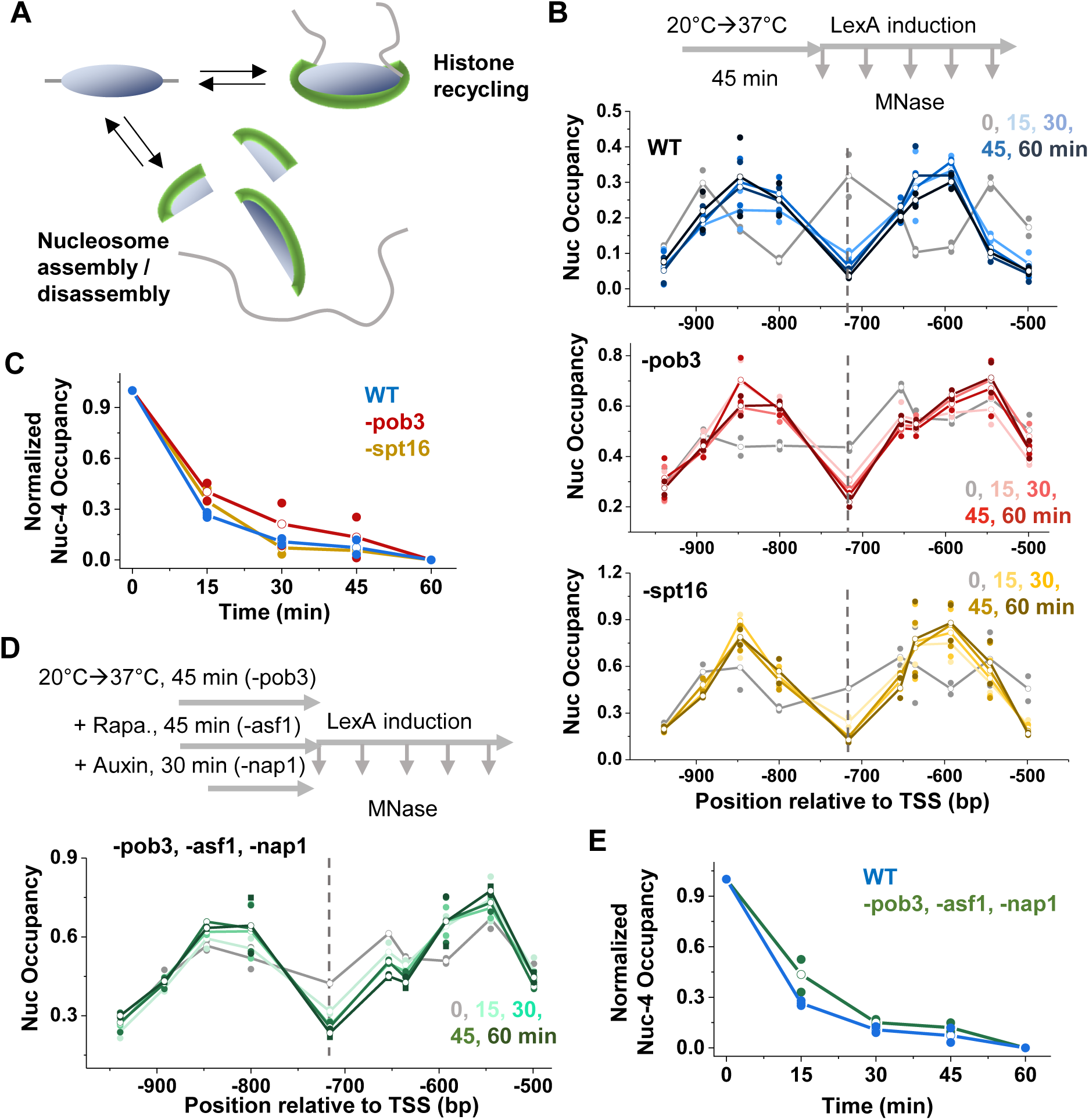
Nucleosome invasion by LexA is not affected by depletion of histone chaperones. **A)** Cartoon illustrating two nucleosome dynamics processes mediated by histone chaperones, histone recycling and nucleosome assembly/disassembly, both of which could enhance the accessibility of nucleosomal DNA. **B)** Time course of LexA invasion into nucleosome –4 in WT (blue), -pob3 (red), or -spt16 (yellow) strains. Pob3 and Spt16 were depleted through temperature-sensitive mutations by shifting cells to the non-permissive temperature of 37°C for 45 minutes prior to LexA induction. **C)** Quantification of nucleosome occupancy change over the LexO site for time course in B. **D)** Time course of LexA invasion into nucleosome -4 with simultaneous depletion of Pob3, Asf1 and Nap1. The three proteins were depleted through temperature sensitive mutation (Pob3), anchor-away (Asf1), and auxin-induced degradation (Nap1). **E)** Quantification of nucleosome occupancy change over the LexO site for time course in D.

The yeast FACT complex consists of two core subunits, Spt16 and Pob3. To probe the relationship between FACT and LexA invasion, we utilized two well-characterized temperature-sensitive mutants, spt16-11 and pob3L78R^38,39^. Shifting these mutant strains from the permissive (22°C) to the nonpermissive temperature (37°C) causes rapid FACT destabilization and degradation^38,39^, ultimately resulting in cell death (**Figure S2A**). We monitored *HOpr* nucleosome occupancy following LexA induction in WT and FACT mutant strains at 37°C. At this elevated temperature, LexA induction occurs faster than at 30°C, leading to more rapid LexA invasion in WT cells (nucleosome –4 is almost completely displaced by 15 min) (**Figure 3B, Figure S2B & C**). Importantly, both Spt16 and Pob3 mutant cells show comparable levels of LexA induction and similar rates of nucleosome –4 displacement (**Figure 3C, Figure S2B & C**).

Given the potential functional redundancy among histone chaperones, we next investigated whether other chaperones compensate for FACT depletion, thereby masking its effect on LexA invasion. To test this, we generated a compound mutant in which the Pob3 mutation was combined with anchor-away depletion of Asf1 (an H3/H4 chaperone) and auxin-induced degradation of Nap1 (an H2A/H2B chaperone) (**Figure S2D & E**). Even in the absence of all three chaperones, LexA can still rapidly and efficiently displace nucleosome –4 (**Figure 3D & E**). This suggests that LexA invasion occurs independently of FACT and other major histone chaperones.

### Rapid replication-independent histone turnover inhibits LexA invasion into nucleosomes

We next tested another dynamic process called histone turnover, where old histones are replaced with new ones. While major histone turnover occurs during DNA replication, it can also take place outside of the S phase through replication-independent mechanisms^40,41^. The impact of cell cycle and DNA replication on LexA invasion will be discussed in the next section; here, we focus on replication-independent histone turnover. Since LexO is engineered at the nucleosome dyad, which predominantly interacts with H3/H4 tetramer, we specifically examined how H3/H4 turnover affects LexA invasion. We hypothesized that higher histone turnover would enhance DNA accessibility at the dyad, thereby promoting LexA invasion (**Figure 4A**).

**Figure 4.**
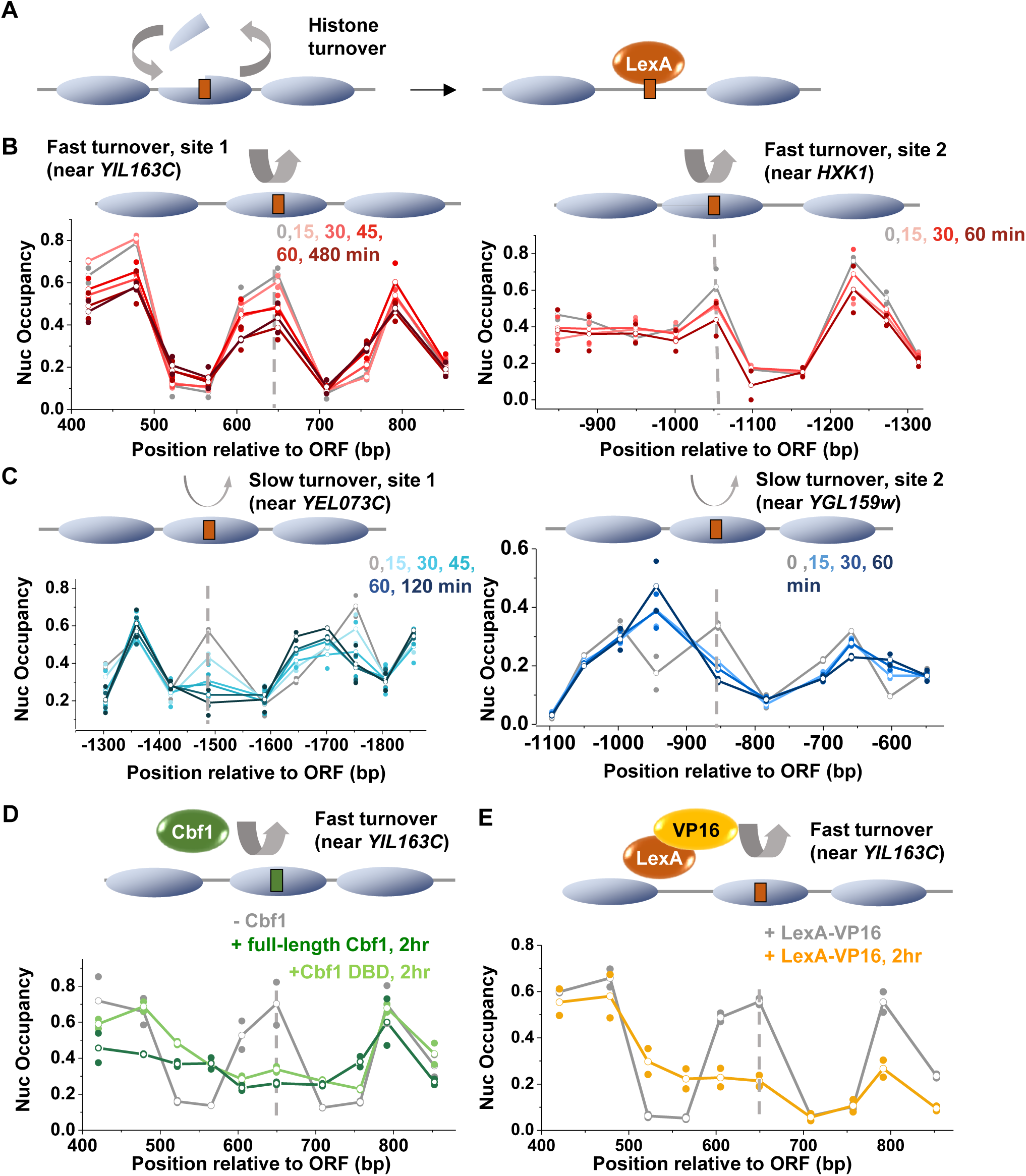
Nucleosome invasion by LexA is inhibited by fast histone turnover. **A)** Cartoon illustrating how fast histone turnover could promote TF binding and nucleosome invasion. **B & C**) Time course of nucleosome occupany near LexO sites engineered into two fast turnover nucleosomes (B) or two low turnover nucleosomes (C). The LexO motifs are positioned near the dyad of these nucleosomes and are marked by the dashed lines. **D)** Same as the left panel in B, except Cbf1 (full length or DBD alone) is induced in a *CBF1* deletion background, with a Cbf1 motif engineered at the same site in the fast turnover nucleosome. Both the Cbf1 full length protein and DBD are able to displace the fast turnover nucleosome. **E)** Same as the left panel in B, except LexA is fused at its C-terminus to the VP16 activation domain. Fusion with VP16 results in nucleosome displacement not observed with LexA alone.

To test this idea, we first examined previously published H3 turnover data in G1-arrested cells^42^. In this dataset, the *HOpr* exhibits a relatively low level of histone turnover. However, the turnover dynamics at our modified *HOpr* construct, which is inserted into a non-native locus, remain unclear. We therefore selected four well-positioned nucleosomes, two with high turnover rates (*YIL163C* and *HXK1*) and two with low turnover rates (*YEL073C* and *YGL159W*) (**Figure S3A & B**), inserted LexO into the dyads of these nucleosomes at their endogenous loci, and measured nucleosome occupancies following LexA induction. Surprisingly, both high-turnover nucleosomes are less displaced by LexA, with their occupancies dropping by only ∼35% after 1 hr of LexA induction, compared to an over 60% reduction at the two low turnover nucleosomes (**Figure 4B & C**).

The unexpected repressive effect of high histone turnover on LexA invasion led us to question whether native yeast PFs like Abf1 or Cbf1 can displace these nucleosomes. In the case of Abf1, we inserted its consensus motif into the same high-turnover nucleosome at *YIL163C* and measured the resulting nucleosome occupancy. Endogenous Abf1 effectively displaces this nucleosome and generates a local NDR (**Figure S3C**), indicating that it has stronger nucleosome displacement activity than LexA. For Cbf1, given our previous finding that the DNA-binding domain (DBD) of Cbf1 confers its high nucleosome affinity^13^, we asked if full-length Cbf1 or the DBD alone can invade into high-turnover nucleosomes. Accordingly, we deleted the endogenous Cbf1 and replaced it with an inducible version encoding either full-length Cbf1 or the DBD alone. Both inducible proteins can invade into the high-turnover nucleosome at *YIL163C* containing engineered Cbf1 motif, although the NDR formed by the DBD is shorter than the one generated by the full-length protein (**Figure 4D**). These findings suggest that strong nucleosome binding by specialized DBDs enhances TF’s capability to open chromatin, while regions outside of the DBD play a role in modulating NDR length, likely through cofactor recruitment.

In yeast, the activation domain (AD) VP16 is known to recruit cofactors such as SAGA and SWI/SNF and facilitate chromatin destabilization and opening^43^. We next asked whether fusing LexA to VP16 could enhance LexA’s ability to invade into high-turnover nucleosomes. Indeed, the LexA-VP16 fusion protein successfully establishes an NDR at the region of high turnover (**Figure 4E**). In summary, these results reveal that rapid histone turnover may present a higher, instead of lower, barrier for TF accessibility, but TF invasion into these regions can be facilitated by strong nucleosome binding via DBD and/or recruitment of co-activators through regulatory domains. Despite these insights, it is still not clear how LexA invades into low-turnover nucleosomes.

### Nucleosome invasion is less efficient during G1 independent of DNA replication

DNA accessibility may vary across the cell-cycle. In particular, DNA replication may transiently expose TF binding motifs behind the replication fork, providing an opportunity for TFs to bind. To investigate the role of cell cycle in LexA-mediated nucleosome displacement, we arrested the cells in G1 with α-factor (**Figure S4A**), induced LexA, and monitored nucleosome occupancy. In G1-arrested cells, nucleosome displacement is significantly slower compared to cycling cells, with the half-life of nucleosome –4 increasing from <30 min to ∼60 min (**Figure 5A & B**). Importantly, LexA expression levels are comparable in G1 vs cycling cells, indicating that the slowed invasion is not due to less LexA induction (**Figure 5C & S4B**).

**Figure 5.**
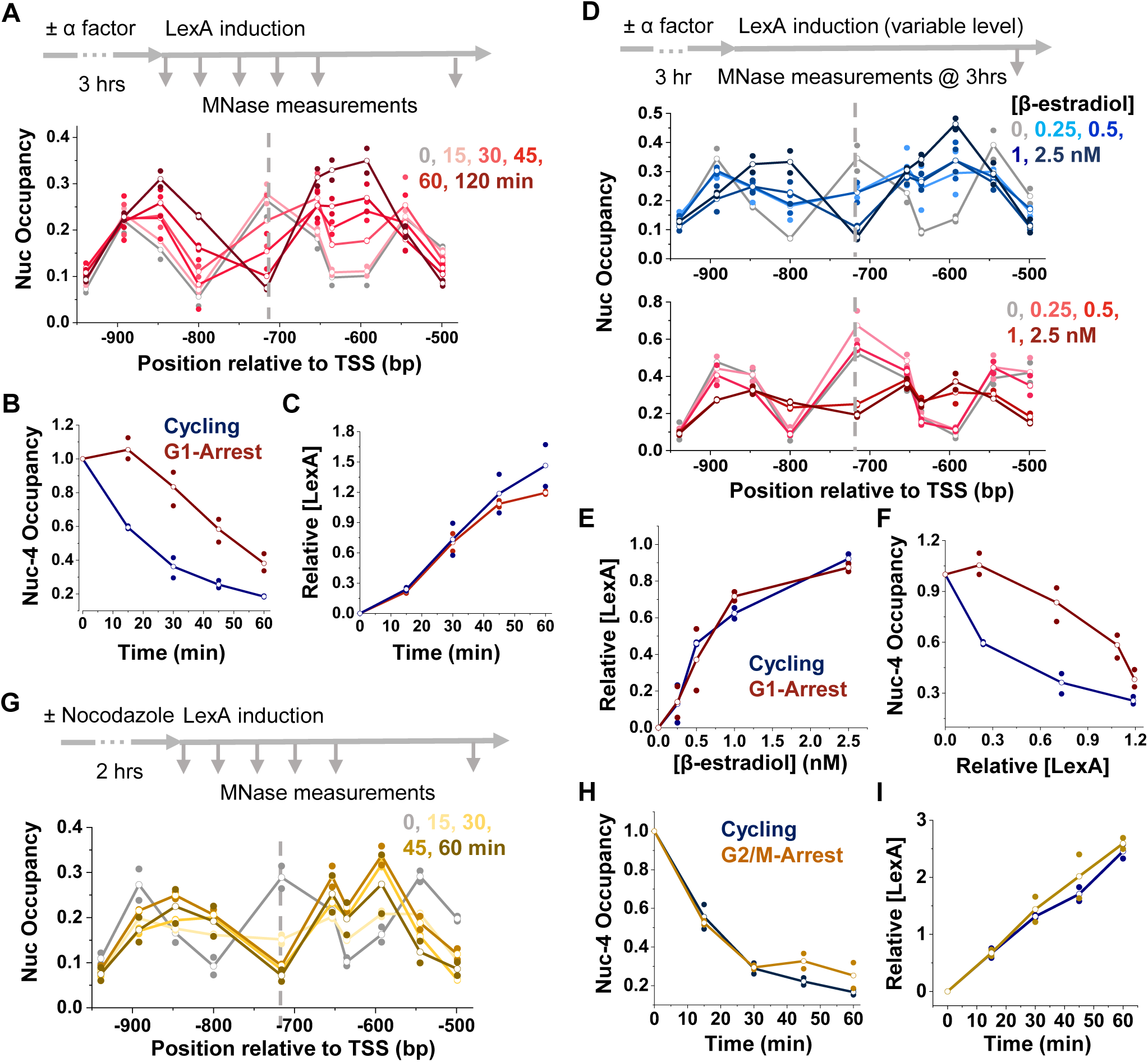
Nucleosome invasion is less efficient during G1, independent of DNA replication. **A)** Time course of LexA invasion into nucleosome –4 in WT cells arrested in G1. **B)** Quantification of nucleosome occupancy at the LexO site during LexA induction in G1-arrested vs cycling cells. **C)** Quantification of LexA levels when induced in G1-arrested (red) vs cycling (blue) cells. **D)** Steady-state nucleosome occupancy near nucleosome –4 following 3 h LexA induction with variable β-estradiol concentrations in cycling (top) or G1-arrested (bottom) cells. **E)** Quantification of LexA levels in response to induction with variable β-estradiol concentrations for G1-arrested (red) and cycling (blue) cells. **F)** Relationship between nucleosome occupancy at the LexO site (from D) and LexA induction levels (from E). **G)** Time course of LexA invasion into nucleosome –4 in WT cells arrested in G2/M by nocodazole. **H & I)** Same as in B & C, but in G2/M arrested vs cycling cells.

Since LexA level is coupled to time during induction, we sought to determine whether the slower nucleosome invasion in G1 was due to a concentration-dependent effect (i.e., a higher level of LexA is required to invade into nucleosomes in G1 and thus takes longer to reach an effective concentration) or a time-dependent effect (i.e., LexA can invade at the same concentration, but the invasion process takes longer in G1). To distinguish between these two possibilities, we induced LexA for 3 hrs to different steady-state levels by titrating 17β-estradiol and measured nucleosome occupancy afterwards (**Figure 5D**). LexA levels were similar between G1-arrested and cycling cells under identical 17β-estradiol concentrations (**Figure 5E & S4C**). Yet, the nucleosome displacement efficiency remains significantly lower in G1 (**Figure 5D**). Combining the data from **Figure 5D & E** shows that higher LexA concentration is required to achieve nucleosome invasion in G1 compared to cycling cells (**Figure 5F**).

To test if the reduced nucleosome invasion in G1 is due to the absence of DNA replication, we arrested the cells in G2/M with nocodazole and repeated the nucleosome occupancy and LexA expression measurements (**Figure 5G-I**). LexA induction levels in G2/M arrested cells are again comparable to cycling cells (**Figure 5I & S4D**). However, unlike in G1, LexA-mediated nucleosome displacement in G2/M occurs at the same rate as in cycling cells (**Figure 5G & H**). Since DNA replication is absent in both G1 and G2/M, yet only G1 showed reduced LexA invasion, these results suggest that the effect is not simply due to the lack of DNA replication. Instead, we suspect that a key factor promoting nucleosome dynamics is downregulated in G1, leading to the observed reduction in LexA-mediated invasion.

### Cohesin facilitates nucleosome invasion by LexA

The observations above prompted us to identify factors that are transcriptionally downregulated in G1 and are also implicated in regulating nucleosome dynamics or chromatin accessibility. Out of the candidates listed in **Figure S5A**, we first investigated the histone acetyltransferase Rtt109, which acetylates newly synthesized H3 at K56^44^. The H3K56ac modification enhances DNA breathing near the nucleosome entry / exit site^45^ and could potentially promote nucleosome dynamics during and after S phase. However, auxin-induced degradation of Rtt109 does not affect nucleosome displacement by LexA in cycling cells (**Figure S5B & C**). We also evaluated Top2, an essential topoisomerase known to relieve DNA torsional strains by resolving both positive and negative supercoils, a process that could influence TF invasion into nucleosomes by affecting nucleosome stability^46,47^. We anchored-away Top2 and confirmed that these cells are no longer viable (**Figure S5D**). Despite this severe phenotype, nucleosome –4 displacement still occurs in the Top2-depleted strain at a rate comparable to control cells (**Figure S5E**).

We next focused on the cohesin complex (**Figure 6A**), an ATPase previously implicated in promoting TF binding and DNA accessibility in higher eukaryotes^48^. All three core cohesin subunits, Smc1, Smc3, and Mcd1, are transcriptionally repressed during G1^49^, and the complex is also actively removed from G1 chromosomes by the cohesin-releasing factors^50^. To test cohesin’s role in nucleosome displacement, we depleted an ATPase subunit Smc3 via auxin-induced degradation (**Figure S6A**) and measured nucleosome occupancy over the *HOpr* following LexA induction in cycling cells. LexA’s invasion rate is already slower in the Smc3-AID strain even without auxin treatment compared to the WT control, consistent with the observed partial degradation of Smc3 in -auxin condition (**Figure S6A & B**). This effect is further enhanced in the presence of auxin, producing a delay comparable to that observed during G1 arrest (**Figure 6B & S6B**). Importantly, LexA induction is not affected by Smc3 depletion, confirming that the observed effect is not due to differences in LexA expression levels (**Figure 6C & S6C**). To further validate these observations, we extended our analysis to other cohesin subunits. Specifically, we targeted Mcd1 (the α-kleisin subunit, known as Rad21 in higher eukaryotes) and Scc2 (the cohesin loader) and repeated the nucleosome occupancy assay. Mcd1 and Scc2 depletion also result in slower LexA-induced nucleosome displacement, although the effects are less pronounced compared to Smc3 depletion (**Figure 6D**).

**Figure 6.**
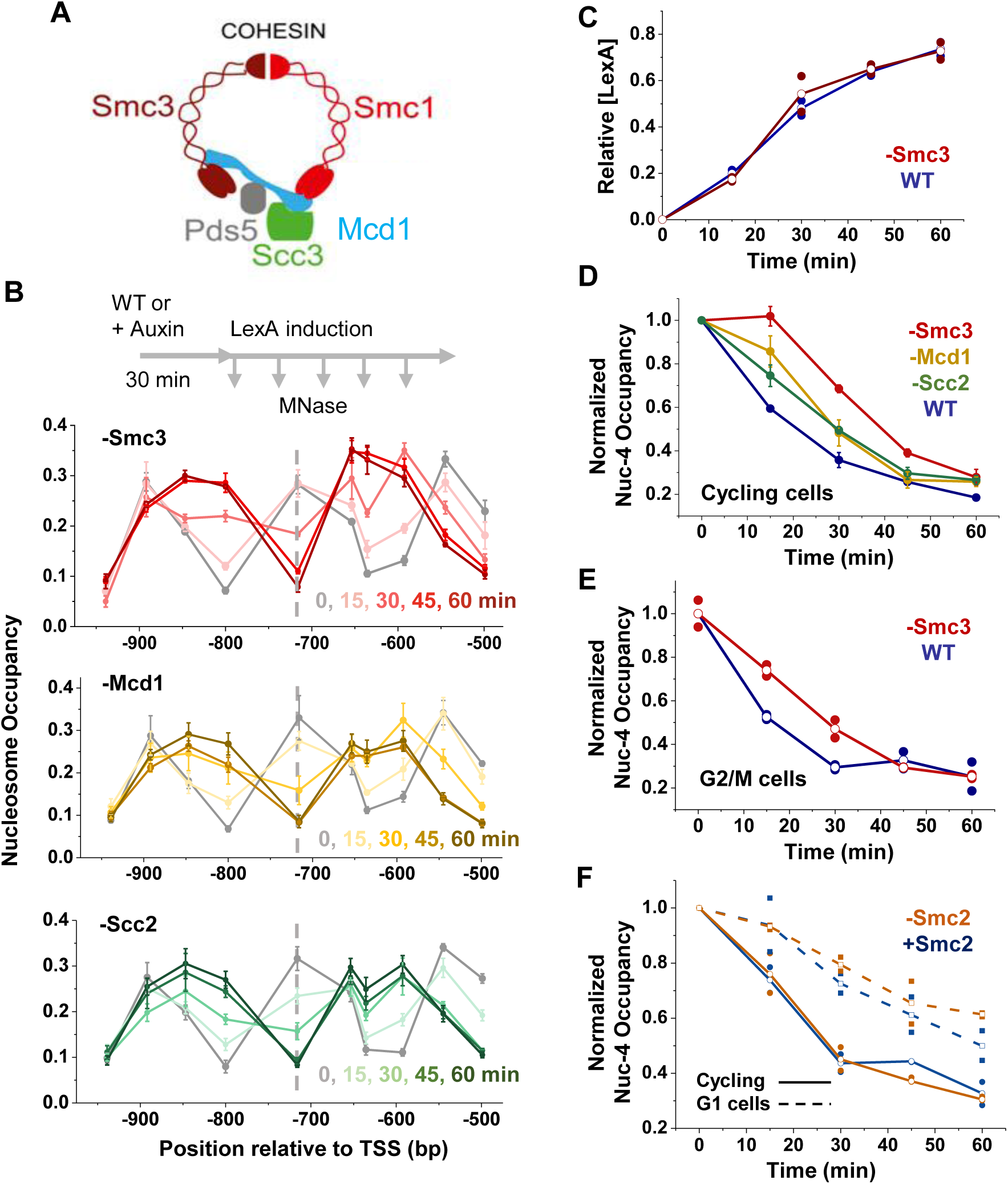
Cohesin facilitates nucleosome invasion by LexA. **A)** Cartoon depiction of the cohesin complex **B)** Time course of LexA invasion into nucleosome –4 in cells depleted of Smc3, Mcd1, or Scc2. **C)** Quantification of LexA expression levels following induction in WT versus Smc3-depleted cells. **D)** Quantification of nucleosome occupancy at the LexO site in WT vs Smc3-, Mcd1-, or Scc2-depleted cells. **E)** Same as in D but measured in G2/M-arrested cells. **F)** Time course of LexA invasion into nucleosome –4 in cells depleted of Smc2, a subunit of the condensin complex.

In human cells, cohesin was proposed to enhance TF binding through a “bookmarking” mechanism, where cohesin maintains chromatin accessibility during mitosis, facilitating TF rebinding in the subsequent G1 phase^48^. This mechanism requires cell cycle progression from M to G1. To test if that’s the case in yeast, we degraded Smc3 in G2/M-arrested cells and measured nucleosome occupancy during LexA induction. Cohesin depletion again resulted in reduced nucleosome displacement (**Figure 6E**), suggesting that the cohesin effect can be decoupled from cell-cycle progression.

Condensin is another SMC complex that shares structural and functional similarities to cohesin. Both complexes have loop extrusion activities, and in yeast, cohesin mainly loop extrudes and condenses chromosome during mitosis, while condensin can loop extrude in G1^51,52^. To test if condensin also contributes to LexA invasion, and specifically, if it promotes LexA-mediated nucleosome displacement in G1, we repeated the LexA induction experiment while depleting Smc2, an ATPase subunit of condensin, in both cycling and G1 arrested cells (**Figure S6D & E**). As observed previously, nucleosome displacement is slower in G1 compared to cycling cells. Smc2 depletion had no detectable effect on nucleosome displacement in cycling cells and caused only a very slight reduction of this rate in G1 (**Figure 6F**), suggesting that condensin plays at most a marginal role in this process. Taken together, these data support a model in which LexA invasion is mainly facilitated by cohesin, with a potential minor role from condensin.

### Cohesin facilitates the binding and nucleosome invasion by a native PF, Cbf1

We next examined whether the effect of cohesin described above extends to a native PF, Cbf1. For this experiment, endogenous Cbf1 was depleted alone or in combination with Smc3 using auxin-induced degradation (**Figure S7A**), and exogenous Cbf1 was reintroduced through 17β-estradiol induction. We then performed MNase and ChIP assays at multiple time points to monitor Cbf1 binding and nucleosome invasion in the presence or absence of cohesin (**Figure 7A**). Western blot analysis confirmed that Smc3 depletion does not affect Cbf1 induction levels (**Figure S7B & C**).

**Figure 7.**
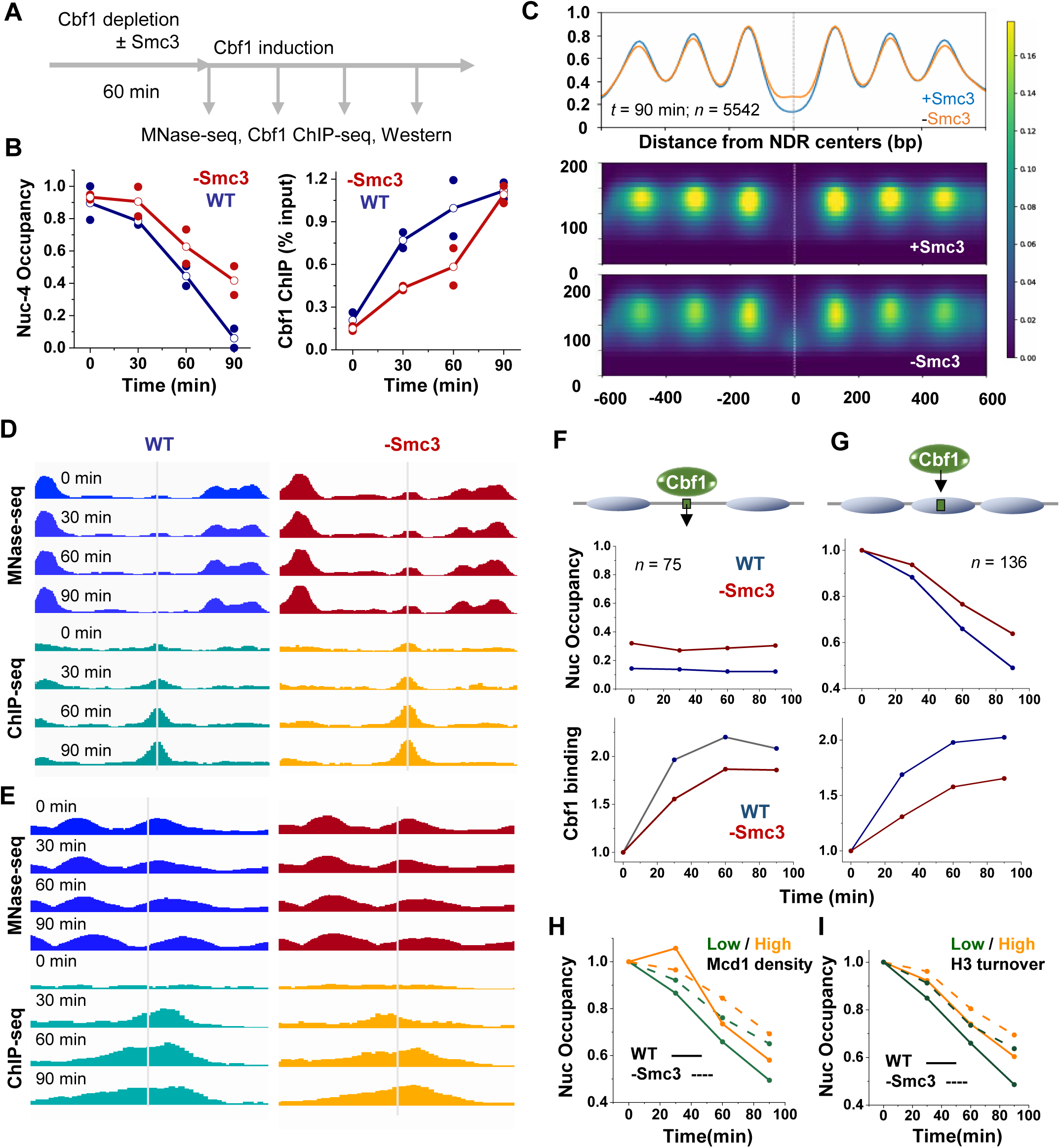
Cohesin facilitates the binding and nucleosome invasion by a native PF, Cbf1. **A)** Experimental workflow for Cbf1 induction ± Smc3 depletion, followed by genome-wide measurement of Cbf1 binding and nucleosome positioning. **B)** Low throughput analysis of Cbf1 binding and nucleosome invasion at an engineered motif positioned near the dyad of *HOpr* nucleosome –4. Left: quantification of nucleosome occupancy at nucleosome –4 following Cbf1 induction in WT versus Smc3-depleted cells. Right: Cbf1 ChIP-qPCR signals at the same site. **C)** Effect of Smc3 depletion on nucleosome occupancy near genome-wide NDRs. Average profile (top) and heatmap (bottom) of nucleosome occupancy aligned at NDR centers. The y axis of the heatmap represents MNase-seq fragment sizes. **D & E)** MNase-seq and Cbf1 ChIP-seq signals over time at two representative loci where Cbf1 binds to open chromatin (D) or within nucleosomes (E) in the presence (left) or absence (right) of Smc3. **F & G)** Average nucleosome occupancy and Cbf1 ChIP-seq signals following Cbf1 induction over all Cbf1 binding sites in open chromatin (F) or embedded within nucleosomes (G). At nucleosome-embedded sites, Cbf1 induction triggers local nucleosome depletion, which is slowed in the absence of Smc3. **H)** Nucleosome depletion at nucleosome-embedded Cbf1 sites during Cbf1 induction, grouped by local cohesin density (based on Mcd1 ChIP-seq signals). **I)** Same as in H but grouped by local H3 turnover signals.

We first carried out a low-throughput analysis using a single engineered Cbf1 motif at the *HOpr* nucleosome –4 dyad. Following Cbf1 induction, nucleosome –4 is displaced and an NDR formed near the Cbf1 motif (**Figure S7D**). Both Cbf1 binding and nucleosome displacement at this site are slowed by Smc3 depletion (**Figure 7B & S7D**), demonstrating that cohesin also facilitates nucleosome invasion by a native PF, at least at this engineered site.

To evaluate this effect at a genome-wide scale, we performed MNase-seq and Cbf1 ChIP-seq during Cbf1 induction. Across all time points (*t* = 0, 30, 60, 90 min), MNase-seq profiles exhibit consistent differences ± Smc3 when aligned at the centers of genome-wide NDRs. Compared to WT, Smc3-depleted cells exhibit higher average occupancy within NDRs and greater fuzziness of flanking nucleosomes (**Figure 7C & S7E**). Since PFs are known to maintain nucleosome depletion within NDRs and form barriers to phase neighboring nucleosomes, these observations suggest that PF activity is compromised in the absence of cohesin.

We next focused on Cbf1 activity at its target sites. ChIP-seq reveals a progressive increase in Cbf1 binding over the induction time course, with 148 ChIP-seq peaks detected at 90 minutes. Most of these peaks (137 out of 148) contain Cbf1 motifs. These motif-containing sites closely match previously mapped endogenous Cbf1 targets, both by ChIP-seq^53^ and ChIP-exo^54^, although the latter identified more sites, likely due to differences in signal-to-noise ratios and thresholding (**Figure S7F**). Among these 137 sites, MNase-seq data at *t* = 0 indicates that 75 reside within broad NDRs (example in **Figure 7D**), 29 are embedded within well-positioned nucleosomes (example in **Figure 7E**), and the rest are in short linkers, at nucleosome edges, or ambiguous (**Figure S7G**). Upon Cbf1 induction, nucleosome-embedded sites underwent progressive nucleosome loss and transitioned into NDRs, consistent with the pioneering activity of Cbf1 (**Figure 7E & S7G**). We also identified 107 additional Cbf1 motifs that show significant nucleosome depletion over time despite lacking strong Cbf1 ChIP-seq signals (**Figure S7G**). These are likely bona fide Cbf1 targets missed by ChIP-seq due to low signal-to-noise or biases against nucleosome-bound events, and most of them overlap with ChIP-exo sites (**Figure S7F**). We therefore combined these 107 sites with the 29 ChIP-seq positive sites as the nucleosome-embedded Cbf1 targets (*n* = 136).

Aggregate analyses of MNase-seq signals centered around nucleosome-embedded Cbf1 sites confirm that nucleosome displacement is slowed down by Smc3 depletion (**Figure 7F & G**). Cbf1 binding is also reduced in the absence of Smc3 for both NDR- and nucleosome-localized sites, with a mildly stronger effect for the latter. For example, after 90 min of Cbf1 induction, Smc3 depletion reduces Cbf1 binding by ∼10% over NDR-localized sites, versus ∼20% over nucleosome-embedded sites (**Figure 7F & G**). Therefore, cohesin facilitates Cbf1 binding to its target sites across the genome, especially at nucleosomal sites.

We next asked whether this effect depends on cohesin’s local enrichment. Nucleosome-embedded Cbf1 sites were stratified by cohesin density based on the Mcd1 ChIP-seq signals (**Figure S7H**)^55^. Surprisingly, sites with low cohesin density show slightly faster nucleosome depletion, and Smc3 depletion slows down Cbf1 nucleosome invasion similarly for both high- and low-density sites (**Figure 7H**). Thus, the effect of cohesin is not limited to sites with high local enrichment and is likely due to broader chromatin changes mediated by cohesin.

Finally, we examined the relationship between Cbf1 invasion and histone H3 turnover. Our data in **Figure 4** shows that Cbf1 can invade into a nucleosome with a high H3 turnover level, but we cannot exclude the possibility that high turnover may affect the dynamics of this invasion quantitatively. To test this, Cbf1 nucleosomal sites were separated into high- and low-turnover groups based on published H3 turnover signals (**Figure S7H**)^42^. Consistent with our findings for LexA (**Figure 4B & C**), Cbf1 displaces nucleosomes with low H3 turnover more efficiently, and both categories show slower nucleosome invasion upon Smc3 depletion (**Figure 7I**). These results indicate that high H3 turnover also has inhibitory effects on Cbf1 binding, and cohesin promotes its nucleosome invasion at sites with both high and low H3 turnover.

## DISCUSSION

By establishing a nucleosome invasion model in WT and mutant budding yeast using the bacterial TF LexA, this study uncovered several surprising findings. First, despite its poor nucleosome-binding ability *in vitro*, LexA can rapidly invade into a well-positioned nucleosome *in vivo* with a binding motif placed at the dyad, an inaccessible region on the nucleosome. Second, the nucleosome invasion capability of LexA does not depend on factors typically associated with nucleosome dynamics, including chromatin remodelers and histone chaperones. Third, contrary to expectations, rapid histone turnover inhibits rather than facilitates nucleosome displacement mediated by LexA and Cbf1. Finally, the cohesin complex, best known for its roles in chromosome compaction and segregation, emerges as a key factor that promotes nucleosome invasion by LexA as well as by endogenous PFs. Although the precise mechanisms underlying these observations remain unclear, we explore some possible explanations below.

### Intrinsic vs context-dependent pioneering activity

PFs are often defined by their ability to bind nucleosomes *in vitro* with similar or only modestly reduced affinities compared to naked DNA. This property is “intrinsic” to these factors, as it arises from direct histone contacts and / or structural features that are compatible with nucleosomal DNA^5,13^. PFs are also functionally defined based on their ability to invade into nucleosomes and open chromatin *in vivo*. The two definitions are generally believed to be linked, i.e. the propensity to bind nucleosomes *in vitro* should facilitate, or even enable, chromatin opening inside cells. However, LexA challenges this view by showing both poor nucleosome binding *in vitro* and efficient nucleosome invasion *in vivo*, clearly demonstrating that the former is not required for the latter. In previous work, we also observed that another bacterial TF, TetR, can bind and displace nucleosomes in yeast, although in that case the motif was placed closer to the nucleosome entry / exit region^16^. These findings suggest that, at least in budding yeast, the ability to invade into nucleosomes may apply to any TFs with sufficiently high DNA-binding affinity and or concentration.

Nonetheless, our data also reveals important distinctions between LexA and native PFs. LexA invasion results in the formation of a very narrow, linker-like NDR, whereas Cbf1 binding at the same position produces a broader NDR (**Figure 1E** vs **S7D**). This difference may be functionally significant, as wider NDRs expose more DNA, potentially allowing additional regulatory factors to be recruited. Furthermore, LexA has difficulty invading into nucleosomes with high histone turnover, while Cbf1, and even its DBD alone, can displace such nucleosomes. These observations suggest that native PFs still possess stronger “pioneering activity” than LexA *in vivo*. Collectively, these observations suggest the need to reconsider how PFs are defined. Rather than solely relying on their nucleosome-binding affinity *in vitro*, defining PFs based on their ability to create broad NDRs or to displace highly dynamic nucleosomes *in vivo* may better reflect their pioneering functions.

### Remodelers function downstream of TF invasion into nucleosomes

By eliminating most remodeling activities in the cells, we demonstrate that CRs are not required for LexA-mediated nucleosome displacement. Importantly, this observation is not unique to the LexA system but is consistent with the behavior of endogenous PFs, as supported by several prior studies. First, major CRs including RSC, ISWI, INO80, and SWI/SNF have been shown to be dispensable for the initial chromatin opening by endogenous PFs, although they play a role in modulating the width of the resulting NDRs^25^. Second, simultaneous depletion of RSC, ISWI, INO80, and CHD1 in yeast has minimal impact on NDRs across the native yeast genome^32^. Third, in live *Drosophila* hemocytes, the chromatin-binding behavior of the PF GAGA factor (GAF) is largely independent of the recruited chromatin remodelers NURF and PBAP, which selectively facilitate the binding of downstream factors^56^.

Together, these results suggest that the primary function of CRs is to shape the size of NDRs following PF binding, rather than to initiate chromatin opening. This idea is supported by recent structural evidence showing that many CRs, including RSC and INO80, bind to extended linker DNA to engage nucleosomes, suggesting that pre-existing NDRs are required for CRs to carry out remodeling activity^57,58^. Thus, a likely sequence of events during NDR formation is that PFs first bind to their target motifs within nucleosomes and generate localized NDRs, which serve as an entry point for CRs to bind and reposition adjacent nucleosomes to further adjust chromatin accessibility.

Notably, SWI/SNF was not depleted in our experiment because its loss severely impairs 17β-estradiol induction^25^, thereby preventing LexA over-expression and subsequent analysis of nucleosome invasion. However, we do not believe SWI/SNF accounts for the residual remodeling activity that enables LexA access. First, SWI/SNF is not very abundant (∼2000 complexes per cell) and thus unlikely to globally remodel the ∼60,000 nucleosomes in the yeast genome to allow rapid LexA invasion into a randomly selected loci. Second, SWI/SNF depletion affects fewer than 100 NDRs genome-wide and does not alter chromatin opening by endogenous PFs^25,59^. Instead, we think that SWI/SNF functions primarily through recruitment, influencing nucleosome dynamics only at the loci to which it is specifically targeted.

### Two strategies to overcome rapid histone exchange

In both yeast and higher eukaryotes, rapid turnover of H3/H4 tetramers predominantly occurs at promoters, enhancers, and within actively transcribed gene bodies^40–42,60^. This dynamic histone exchange is usually thought to be associated with transcriptional activation, as it disrupts the canonical nucleosome structure and transiently exposes otherwise occluded DNA. Contrary to this view, we found that LexA is less efficient at invading into nucleosomes with rapid histone turnover, suggesting that at least some turnover sites may inhibit, rather than enhance, DNA accessibility.

The underlying mechanism for this inhibitory effect remains to be elucidated. However, the observation that the same high-turnover nucleosome can be displaced by endogenous PFs, including the Cbf1 DBD, and LexA fused with VP16 raises the following possibilities. Regions with rapid histone turnover are sites where nucleosomes are continuously disassembled and reassembled, and newly deposited histones may form structures that disfavor LexA binding, given its inherently low affinity and fast dissociation from nucleosomal DNA. In contrast, Cbf1 or its DBD may remain stably bound during this disruptive process due to its ability to engage nucleosomal DNA. On the other hand, LexA-VP16 may persist at these sites by recruiting activating co-factors, such as SWI/SNF and SAGA, which can destabilize and disrupt nucleosomes and clear the roadblock for LexA to stably bind.

These findings support the existence of two mechanistically distinct strategies by which TFs can achieve stronger pioneering activity *in vivo*. The first is to develop specialized DBDs that can better dock onto nucleosomes, like what was observed for yeast PFs Reb1 and Cbf1^11,13^ and several PFs in higher eukaryotes^5,6,61^. Alternatively, TFs can use regions outside of DBD to recruit co-factors to enhance chromatin interaction and overcome the nucleosome barrier. Note that these two mechanisms are not mutually exclusive. Some PFs like FOXA1, SOX2, and Zelda may indeed employ both strategies for their chromatin opening and stable occupancy^62,63^.

### Promotion of PF activity by the cohesin complex

By depleting cohesin subunits, we demonstrate that the cohesin complex enhances chromatin accessibility at genome-wide NDRs and facilitates nucleosome invasion by both the artificial TF LexA and the endogenous PF Cbf1. These results are consistent with a previous study in mammalian cells showing that cohesin is broadly associated with TF binding clusters, and that cohesin loss leads to reduced DNA accessibility and TF occupancy^48,64^. Together, these observations suggest that cohesin may play a conserved role in promoting PF activity.

A previous mammalian study attributed this effect to a “mitotic bookmarking” mechanism, whereby cohesin remains bound during early mitosis to maintain open cluster sites, enabling TFs to re-engage in late M / next G1 phase^48^. However, this model does not apply to budding yeast, where TFs are not broadly displaced during M phase due to modest chromatin compaction^65^. Consistently, LexA induced at G2/M can engage chromatin and mediate nucleosome displacement (**Figure 4D**). In addition, the bookmarking mechanism requires M-G1 transition, yet the effect of cohesin on LexA invasion can be detected in cells arrested in G2/M without cell cycle progression. Therefore, the effect of cohesin on chromatin accessibility and TF binding is unlikely due to bookmarking in yeast.

Although the precise mechanism remains unclear, two observations provide important clues. First, cohesin promotes Cbf1 binding at both NDR-localized and nucleosome-embedded sites. Second, this effect is not restricted to loci with strong cohesin enrichment. These results argue against a simple “site holding” model in which cohesin directly maintains open chromatin at its binding sites. Instead, cohesin appears to exert a global influence on chromatin accessibility. One possibility is that the passage of cohesin during loop extrusion physically reorganizes or untangles chromatin fibers, thereby enhancing accessibility^66^. In addition, cohesin activity has been shown to generate torsional stress on chromatin^67^, which can modulate the binding specificity and stability of nucleosomes and TFs^46^. Further experiments will be required to test these hypotheses.

## RESOURCE AVAILABILITY

### Lead contact

Further information and requests for resources and reagents should be directed to and will be fulfilled by the lead contact, Lu Bai.

### Materials availability

Requests for yeast strains should be directed to Lu Bai. This study did not generate any new reagents.

### Data and code availability

- MNase and ChIP seq data have been deposited at GEO at GEO: GSE311346 and GSE311347 and are publicly available as of the date of publication. Previously published data were used in this study are available at GEO: GSE1433 (turnover chip-seq) GSE1514 (cohesin chip-seq) GSE252386 (Cbf1 chip-seq) and GSE147927 (Cbf1 ChIP-ExO).
- Any additional information required to reanalyze the data reported in this paper is available from the lead contact upon request.

## ACKNOWLEDGMENTS

We thank Dr. Joseph Reese for providing anchor-away plasmids and yeast strains. We thank Dr. David Stillman for providing temperature-sensitive FACT yeast strains. We thank Dr. David Shore for providing a background chromatin-remodeler degradation strain. We acknowledge the Genomic Research Incubator at Penn State (Cheryl Keller) and Huck Institutes’ Flow Cytometry Core Facility for technical support. We acknowledge all members in the Bai lab and Poirier lab for insightful comments on the manuscript. We also thank the members of the Center of Eukaryotic Gene Regulation at Pennsylvania State University for discussions. This work is supported by the National Institutes of Health (R35 GM139654 to L.B. and R35 GM139564 to M.G.P.).

## AUTHOR CONTRIBUTIONS

S.S., H.C. and L.B. designed the experiments; S.S. performed most of the yeast experiments and data analysis; H.C., C.S., S.V., and S.B. contributed to the yeast experiments; R.C. and M.P. conducted the *in vitro* EMSA experiment; S.S. and L.B. wrote the manuscript.

## DECLARATION OF GENERATIVE AI AND AI-ASSISTED TECHNOLOGIES

During the preparation of this work, the author(s) used ChatGPT to polish the writing of the manuscript. After using this tool or service, the author(s) reviewed and edited the content as needed and take(s) full responsibility for the content of the publication.

## SUPPLEMENTAL INFORMATION

**Document S1. Figures S1–S7 and Table S1.**

**Table S1. Primers and strains.**

**Document S2. Article plus supplemental information.**

## STARfllMETHODS

### KEY RESOURCES TABLE

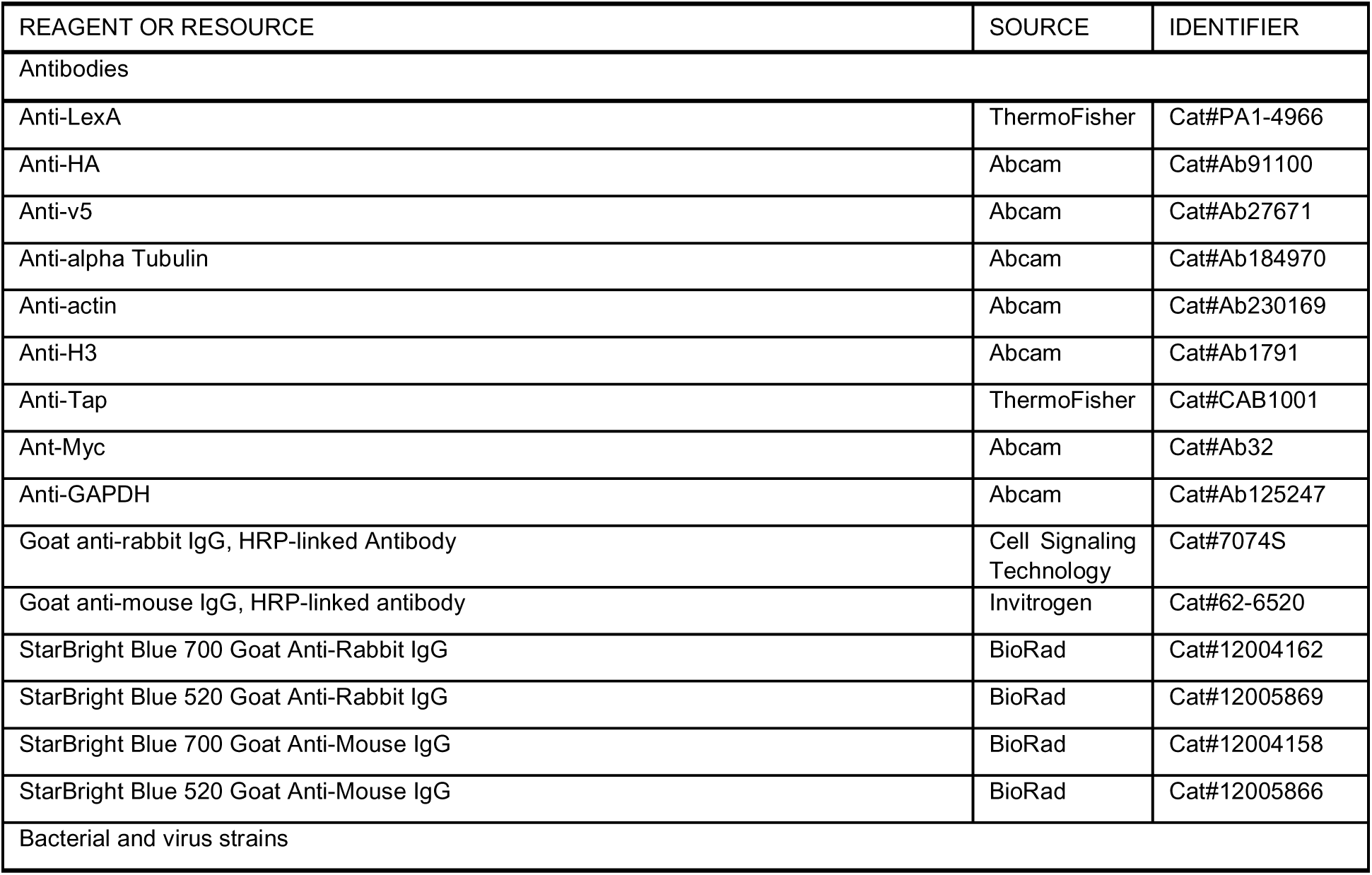
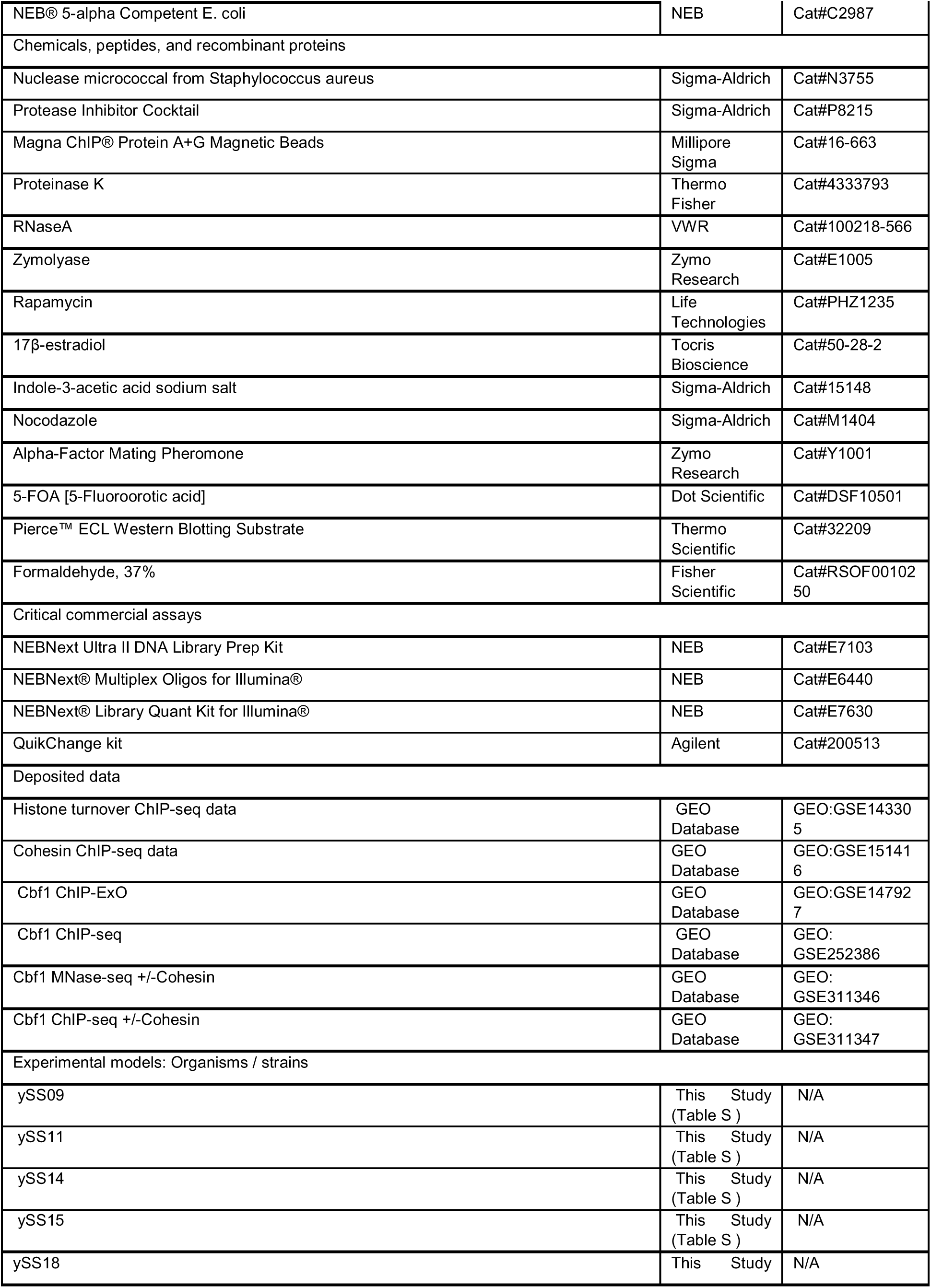
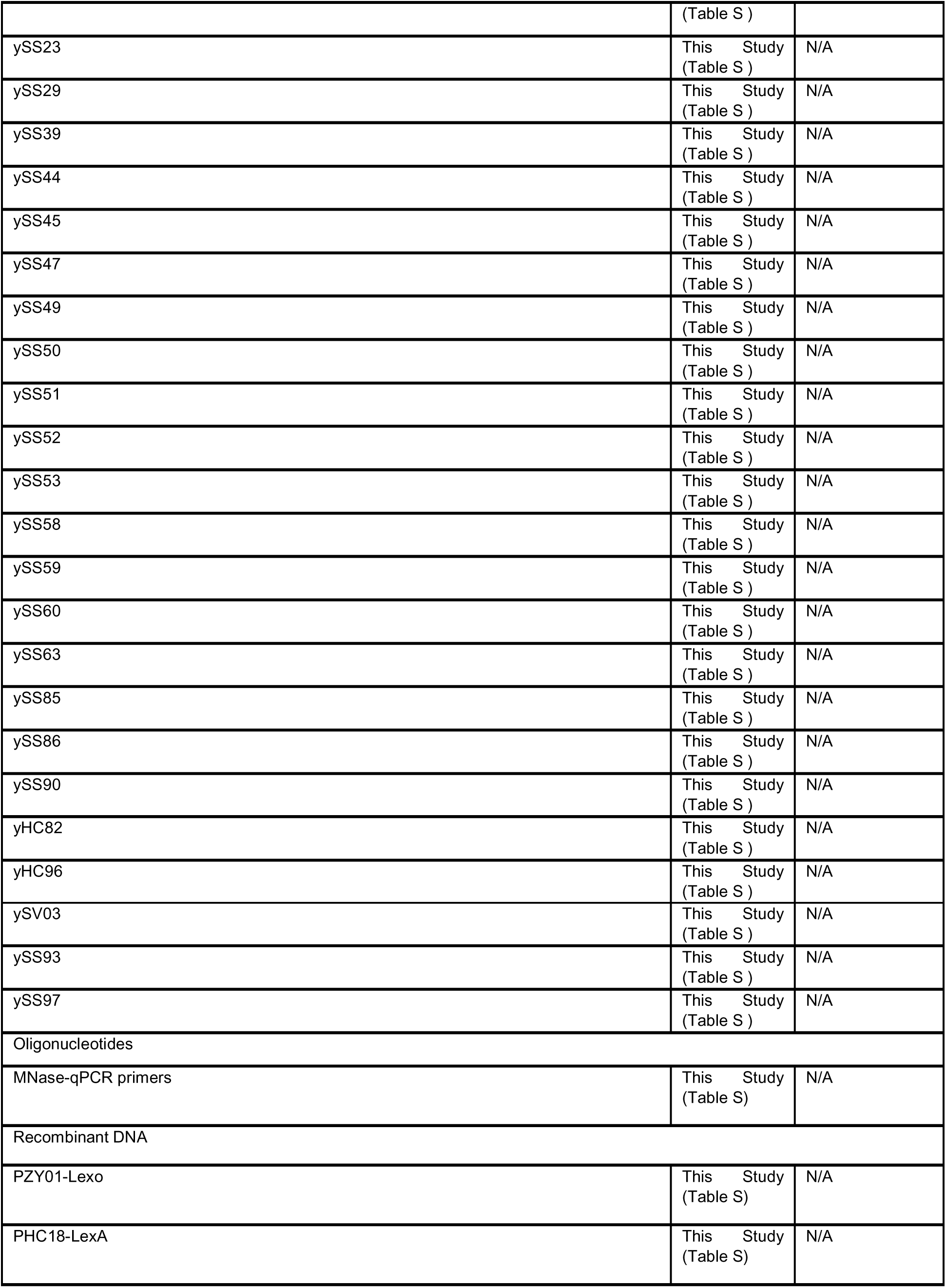
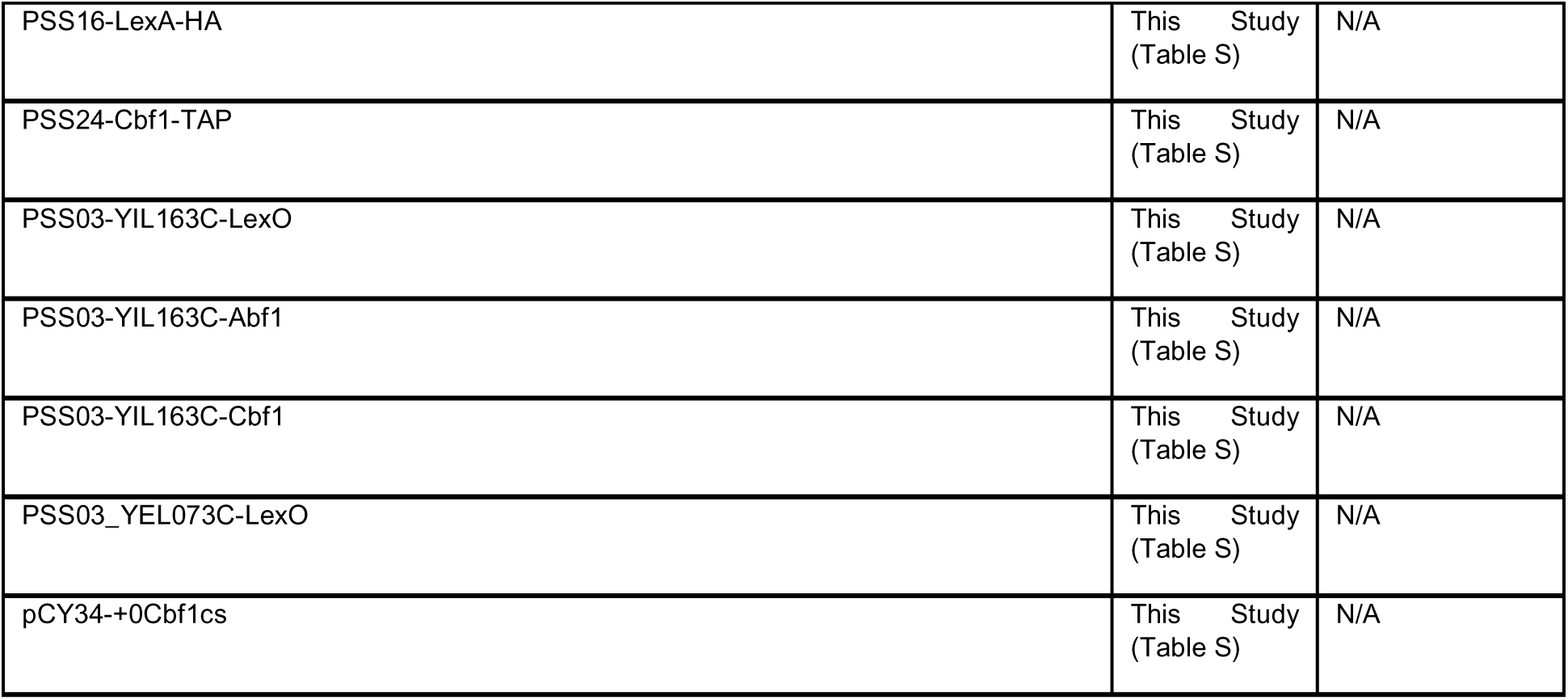

### EXPERIMENTAL MODEL AND STUDY PARTICIPANT DETAILS

#### METHOD DETAILS

##### Plasmid and yeast strains

Standard methods were used for strain construction and plasmid cloning. All yeast strains were derived from w303 background. The endogenous *HO* promoter (−1298 to −272 relative to *HO* ORF) was deleted, and a modified *HOpr* lacking Swi5 binding sites was inserted into the *CLN2* locus^16^. LexA or LexA-HA, driven by a core *GAL1* promoter, was integrated at the *HIS3* locus. The same strain also expressed Gal4-ER-VP16, allowing LexA induction in the presence of 17β-estradiol. Engineering LexO / Cbf1 / Abf1 motifs into loci with various level of H3 turnover was done using an FOA pop-out scheme with seamless editing.

For auxin induced degradation strains, a 3x-v5-AID DNA cassette was amplified with primers containing 70 bp of DNA homology arms and integrated by homologous recombination to generate in-frame gene fusions. Protein degradation was verified by treating the samples with 500 µM 3-indoleacetic acid (IAA) (Sigma, I2886), typically for 30-60 min, followed by western blotting. Anchor-away strains were generated by tagging the target with FRB-TAP-GFP. Nuclear protein depletion was induced by treatment with 1 μg/mL rapamycin for 45 min and confirmed by spotting assays or fluorescence microscopy. Temperature-sensitive FACT mutants were grown at 20°C for all experiments unless otherwise indicated. For inactivation, cells were shifted to 37°C for 45 min.

For the inducible Cbf1-TAP strain in Figure 3, we replaced the endogenous *CBF1* promoter with *GAL1* promoter. An inducible Cbf1-TAP driven by *MET3* promoter was integrated at the *HIS3* locus. Cells were first grown in SCD+20x Met to repress the endogenous Cbf1 expression and shifted to SCD-met to induce Cbf1-TAP. The Cbf1-TAP strain in Figure 7 was generated by tagging the c-terminus of endogenous Cbf1 with an AID-tag. An inducible Cbf1-TAP driven by a *GAL1pr* was inserted in the *HIS3* locus which can be induced with 17β-estradiol.

For cell-cycle arrest experiments, G1 arrest was achieved with 5 µM α-factor (Zymo Y1001) in SCD-Met medium for 3 h (equivalent to the doubling time of the anchor-away strain in synthetic media)^68^. G2/M arrest was achieved with 15 μg/mL nocodazole (Sigma M1404) for 2 h (one doubling time in YPD).

##### Cell cycle analysis

Cycling or arrested cells were grown to an OD of 0.4, and 1 mL culture from each condition was collected for FACS analysis^69^. Cells were fixed with ethanol overnight and stained with 2.5 µM Sytox Green in 50 mM sodium citrate buffer containing 20 µg/mL RNase A. Following overnight proteinase K treatment (10 µL, 20 mg/mL), the cells were pulse-sonicated to reduce aggregates. FACS analysis was carried out using a Cytek Aurora (Cytek Biosciences) analyzer at the Huck Flow Cytometry Facility. The instrument was configured using conventional mode and gain settings were optimized for FSC, SSC and Sytox Green signals. Single cells were gated using FSC-A/H and Sytox Green-A/H bivariant plots. The DNA intensity (Sytox Green) was displayed in a histogram using linear scale with the G1 peak near the first quartile and the G2 peak near the second quartile of the histogram axis. For each sample 50,000 cells were recorded, and the data files were analyzed using Kaluza software (Beckman Coulter).

##### *In vitro* nucleosome assembly

The YNN1 sequence was cloned in the pUC19 plasmid. A LexA-binding site (TACTGTATGAGCATACAGTA) was introduced at the p65 position (65 bp from the entry/exit site centered at the dyad) by site-directed mutagenesis (QuikChange kit, Agilent). DNA templates were labeled at the 5’ end with Cy3 using labeled oligonucleotides from IDT. All DNA constructs were amplified by PCR and purified using an anion-exchange MonoQ column (Cytiva) on an HPLC system (Agilent). Purified DNA was stored in 0.5X TE buffer (5 mM Tris base, 0.5 mM EDTA).

Recombinant histones (human H2A with K119C, human H2B, *Xenopus laevis* H3 with C110A, and human H4) were purchased from “The Histone Source” (Colorado State University; https://histonesource-colostate.nbsstore.net/). Histone octamers were refolded under a 1:1.3 molar ratio of H3 and H4 to H2A and H2B. Site-specific labeling was performed by conjugating Cy5-maleimide to H2A K119C. Labeled octamers were purified on a Superdex 200 column (Cytiva) using an ÄKTA Pure FPLC system (Cytiva). Nucleosomes were reconstituted by salt dialysis using purified DNA and histone octamers mixed at a 1.5:1 molar ratio. The reconstituted nucleosomes were purified using sucrose gradient centrifugation on an Optima L-90K Ultracentrifuge (Beckman) at 41000 rpm under 4 □C for 22 hrs. The 5%-30% sucrose gradient was prepared with 0.5x TE buffer using Gradient Master (Biocomp). Purified nucleosomes were stored in 0.5x TE with 20% glycerol and aliquoted in −80 □C.

##### Electrophoresis mobility shift assay (EMSA)

0.5 nM DNA or nucleosomes were incubated with variable concentrations of LexA in T130 buffer (10 mM Tris-HCl pH 8.0, 130 mM NaCl, 10% glycerol, and 0.0075% Tween-20) under room temperature for 20 minutes. The mixtures were run on a 5% acrylamide native gel with 0.3x TBE buffer (30 mM Tris base, 27 mM boric acid, and 0.3 mM EDTA) for 90 minutes at 300 V.

##### MNase assay and MNase-seq

1.5 OD_660_ units of cells were harvested and washed sequentially with 1 mL water and 1 mL 1M sorbitol. The pellet was re-suspended in 0.5mL spheroplasting solution (1 M sorbitol, 0.5 mM 2-mercaptoethanol, 0.18 mg/ml zymolyase) and incubated at room temperature for 7 minutes with gentle inversion. Spheroplasts were collected by centrifugation at 500 x g for 3 min, washed twice with 1 mL of 1 M sorbitol, and resuspended in 200 μL digestion buffer (1 M sorbitol, 50 mM NaCl, 100 mM Tris-Cl pH 7.4, 5 mM MgCl_2_, 1 mM CaCl_2_, 1 mM 2-mercaptoethanol, 0.5 mM spermidine, 0.075% NP-40). Samples were incubated at 37°C for 8 min with 1U of MNase (Sigma N3755). Reactions were terminated by adding 20 μL quench buffer (250 mM EDTA and 5% SDS). DNA was purified by phenol–chloroform extraction, and mono-nucleosome-sized DNA was gel-extracted before proceeding to stacking qPCR^70^. Nucleosome occupancy was normalized to a reference nucleosome in the *EXO84* terminator^25^.

For MNase-seq, 1.5 OD_660_ units of cells were digested with 1U (Sigma N3755) of MNase for 8 minutes which resulted in ∼80% genome converted into mono-nucleosomes. 100 ng of MNase-digested DNA was used to generate sequencing libraries using NEBNext® Ultra™ II DNA Library Prep Kit for Illumina® (Catalog # E7645L), following the manufacturer’s protocol. Libraries were sequenced on an Illumina NextSeq 2000 in 2 × 50 bp paired-end mode. ∼20 million reads per sample were generated and aligned to the sacCer3 genome assembly using Bowtie2 with presets “--very-sensitive”. Bigwig files were generated using deepTools bamcoverage (version 3.5.6) using a 130-170bp size selection and alignments were trimmed by 15bp on either side using offset 15 −15. Two biological replica were performed at each time point.

##### Western blotting and quantification

Yeast cultures (10 mL) were grown to ∼0.15 OD_660_, harvested, washed with 0.5 mL H_2_0, and resuspended in 200 μL 1 M NaOH. After incubation at room temperature for 10 min, cells were pelleted and resuspended in 50 μL SDS–PAGE sample buffer. Proteins were resolved by SDS–PAGE and transferred for western blotting using standard protocols. Primary antibodies were used at the following concentrations: anti-LexA (ThermoFisher,PA1-4966) 1:3000, anti-HA (abcam, ab91100) 1:5000, anti v5 (abcam, ab27671) 1:2000, anti-alpha tubulin (abcam, ab184970) 1:15000, anti-actin (abcam, ab230169) 1:2000, anti-H3 (abcam, ab1791) 1:2000, anti-GAPDH (abcam ab125247) 1:5000, anti-TAP (ThermoFisher, CAB1001) 1:1000, anti-myc (abcam, ab32) 1:1000. Secondary antibodies were used at the following concentrations: goat anti-rabbit igG (Cell Signaling Technology, 7074s) 1:3000, goat anti-mouse IgG (Invitrogen, 62-6520) 1:2000, goat anti-rabbit (Biorad, 12004162 & 12005869) 1:3000, goat anti-mouse (Biorad, 12004158 & 12005866) 1:3000.

To enable direct comparisons, inducible LexA and endogenous Cbf1, Reb1, and Ser2 were tagged at their C-terminal with an HA tag. Reb1 and Cbf1 are known budding yeast PFs and serve as a good internal comparison for monitoring the expression of LexA. Protein copy numbers were obtained from SGD (Cbf1, 5757 +- 2714, Reb1, 5582 +- 1829, and Ser2, 11714 +- 2168) and converted to concentrations using the average volume of 42 μm^3^ and nuclear volume of 3 μm^3^ in haploid yeast ^71^. The western blot intensities of these proteins agreed well with their known copy numbers within the cell (**Figure S1C**).

For quantification, four concentrations of lysate were analyzed per sample to ensure signals were within the linear detection range (only data within the linear range were used). A loading control confirmed equal protein loading, and an internal reference standard was included for normalization across blots. For Cbf1-TAP quantifications in **Figure S7B**, normalization was performed against total protein content combined with an internal standard. This is because the usual loading control, GAPDH, is regulated by Cbf1 (based on previously published RNA-seq measurements^72^), and its expression varied in Cbf1-AID strains ± auxin.

##### ChIP-seq

Yeast cultures (50 mL) were grown in SCD–Met medium to an OD_660_ of 0.4. Crosslinking was performed by adding 1.39 mL of 37% formaldehyde directly to each culture and incubating at room temperature for 20 min. Crosslinking was quenched with 2.7 mL of 2.5 M glycine for 5 min. Cells were pelleted at 3000 rpm for 3 min at 4°C, resuspended in 1 mL TBS, and washed twice with 1 mL TBS. Cells were lysed by resuspension in 250 μL FSPP buffer (50 mM HEPES-KOH pH 7.5, 140 mM NaCl, 1 mM EDTA, 1% Triton X-100, 0.1% sodium deoxycholate, 1% protease inhibitor cocktail, 1 mM PMSF) with 300 μL glass beads. Samples were vortexed at 4°C twice for 20 min each, separated by a 10 min break. Lysates were released by piercing the tube ends with a hot needle and centrifuging at 3000 rpm for 5 min at 4°C into 15 mL tubes. Lysates were recovered, supplemented with 1 mL FSPP buffer, and sonicated (30 s on/off cycles for 7 min). After centrifugation at 14,000 rpm for 20 min at 4°C, 200 μL supernatant was saved as input control.

For immunoprecipitation, 200 μL of lysate was diluted with 800 μL FSPP buffer and incubated with 20 μL pre-blocked magnetic IgG beads and 1–2 μg of antibody (TAP: Thermo CAB1001; LexA: ThermoFisher PA1-4966; V5: Abcam ab27671) overnight at 4°C with rotation. Beads were sequentially washed with 1) FA Lysis buffer (50 mM HEPES-KOH pH 7.5, 140 mM NaCl, 1 mM EDTA, 1% Triton X-100, 0.1% sodium deoxycholate), 2) FA Lysis + 150 mM NaCl, 3) FA Lysis + 500 mM NaCl, 4) LiCl buffer (0.25 M LiCl, 1% NP-40, 1% sodium deoxycholate, 1 mM EDTA, 10 mM Tris-HCl pH 8.0), and 5) TE buffer (pH 8.0). ChIP sample was then eluted with 400 μL elution buffer (50mM NaCl, 50mM Tris-HCl, pH 8.0, 10mM EDTA, 1% SDS) by rotating at 30°C for 30 min. Eluates were treated with 5 μL of 20 mg/mL proteinase K overnight at 65°C. Input controls were processed in parallel by adding 160 μL FA Lysis buffer, 40 μL 10% SDS, and 5 μL proteinase K, followed by overnight incubation at 65°C.

DNA was extracted with phenol:chloroform:isoamyl alcohol. ChIP samples were resuspended in 40 μL H_2_0. Input samples were resuspended in 380 μL TE buffer with 10 μL of 10mg/mL RNase A and incubated at 37°C for 30 minutes. DNA was washed with EtOH and resuspended in 40 μL H_2_0. Factor enrichment was verified through qPCR. For ChIP-seq, ∼5 million paired-end reads were generated and aligned to *S. cerevisiae* genome sacCer3 with Bowtie2 (version 2.5.4) using default parameters. Peak calling was performed with MACS2 (version 2.2.7.1). Bigwig files were generated using deepTools bamcoverage (version 3.5.6). Two biological replica were performed at each time point.

#### QUANTIFICATION AND STATISTICAL ANALYSIS

For paired-end MNase-seq data, dyad and nucleosome positions were inferred using DANPOS3, which calculates fragment midpoints and genome-wide dyad density^73^. In addition to average nucleosome coverage from all mapped reads (**Figure S7E**), we generated V-plots^74^ to provide more information about nucleosome distributions by sorting the sequencing reads according to their fragment length (**Figure 7C**).

For Cbf1 analysis, we first scanned the yeast genome for potential consensus binding sites using published position weight matrix and the recommended cutoff^75^. For each consensus site, we quantified both the MNase-seq signal at the motif center and the Cbf1 ChIP-seq signal within ±100 bp of the motif. Across the Cbf1 induction time course, nucleosome occupancy and Cbf1 enrichment at each site were normalized to the *t* = 0 condition, such that these values represent a fold change relative to baseline. Based on the combined ChIP-seq and MNase-seq profiles, we classified Cbf1 binding sites into four categories: 1) ChIP+ & NDR-localized, 2) ChIP+ & nucleosome-embedded, 3) ChIP- & nucleosome-embedded, and 4) nucleosome edge or ambiguous (**Figure S7G**). Similarly, ChIP-seq densities for cohesin and H3 turnover were determined for each Cbf1 binding site using previously published datasets^42,55^ and grouped according to their density.

---

**Figure S1.**
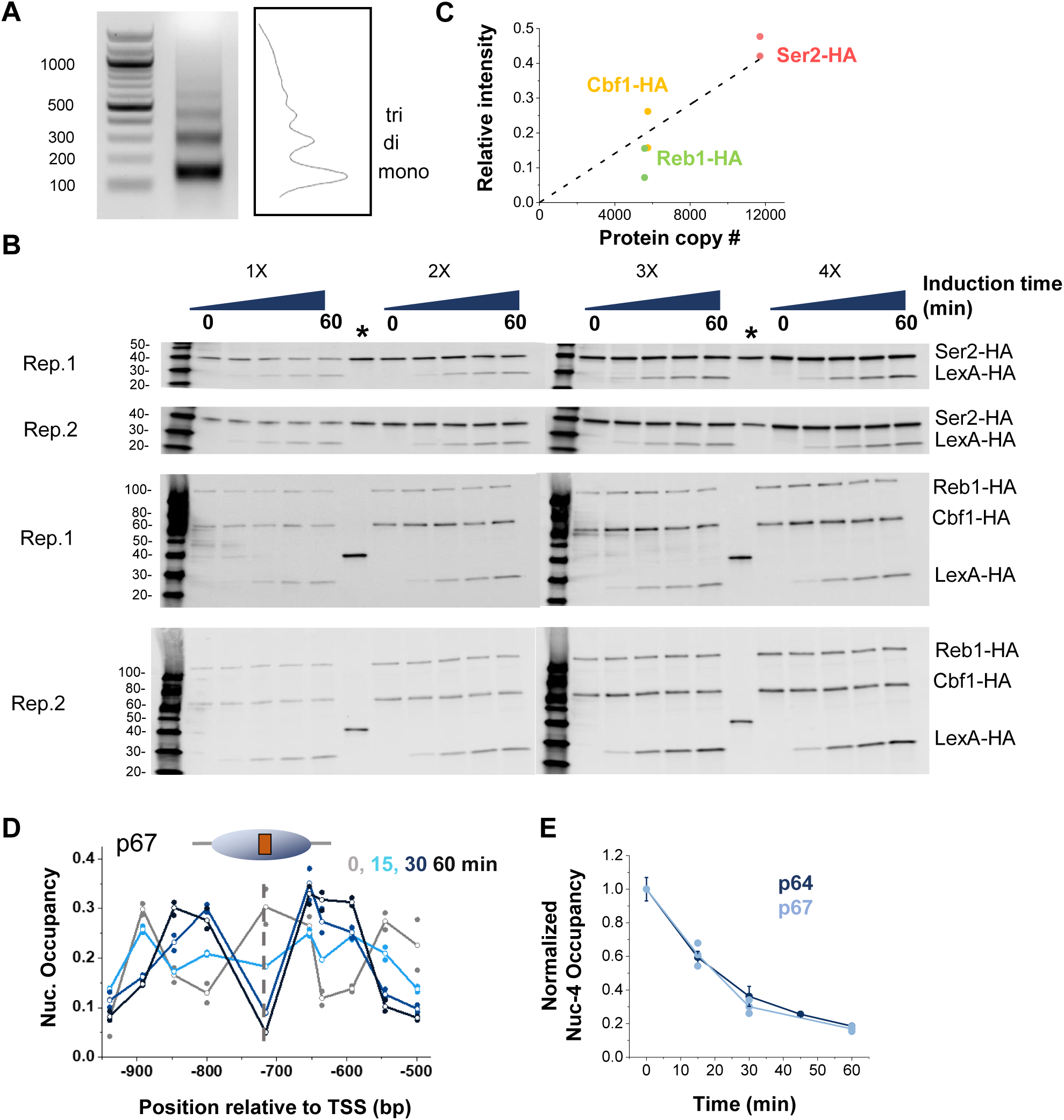
Additional LexA invasion and quantification data, related to Figure 1. **A)** Representative MNase gel and the corresponding ImageJ intensity plot showing mono-, di-, and tri-nucleosome bands. Mononucleosome bands were gel extracted for nucleosome mapping by qPCR. **B)** Western blots (two biological replicates for each time point and strain) used for quantifying LexA-HA levels in Figure 1H. Four different cell lyaste concentrations were loaded ensure that each time point fell within the linear detection range of the assay. The asterisks mark the gel lanes containing internal HA loading control (see Methods). Molecular weight markers indicated in first lane. **C)** Band intensity vs the known copy number of Cbf1, Reb1, and Ser2 (from SGD). The linear fit (gray curve) was used to estimate the copy number of LexA. **D)** Time course of LexA invasion with a slightly shifted LexO position in nucleosome –4. The original dyad position of LexO in Figure 1D is p64, here shifted by 3 bp to p67. **E)** Quantification of nucleosome depletion during LexA induction over the LexO site at p64 and p67.

**Figure S2.**
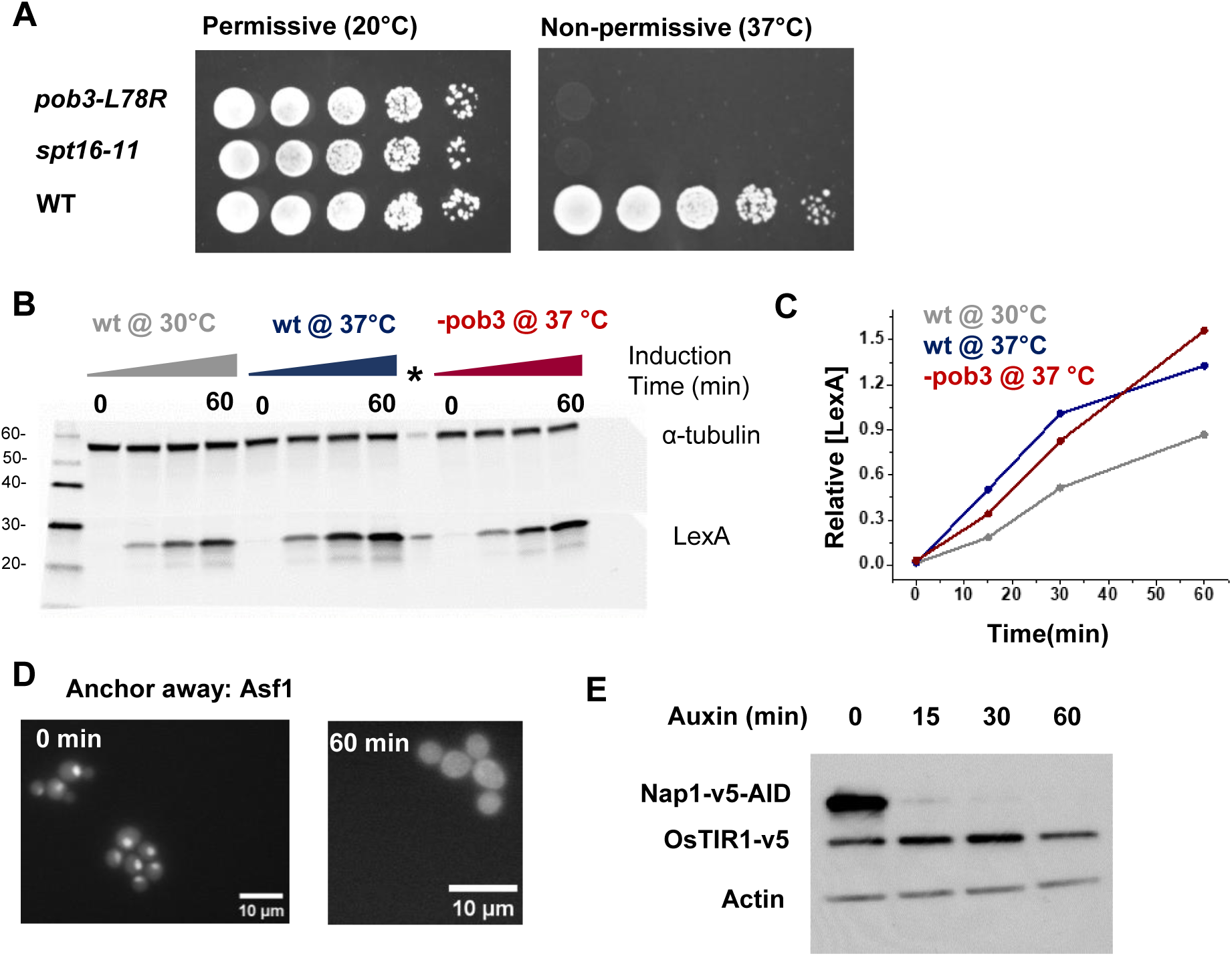
Histone chaperone depletion and LexA induction, related to Figure 3. **A)** Spotting-assay for WT yeast and two temperature-sensitive FACT mutants grown at permissive (20°C) and nonpermissive (37°C) temperatures. **B)** Western blot analysis of LexA induction in WT and FACT mutant strains. Molecular weights are indicated in the first lane. **C)** Quantification of LexA levels in the Western blot in B. **D)** Fluorescence images showing anchor-away of Asf1-FRB-GFP following rapamycin treatment. **E)** Western blot confirming auxin-induced degradation of Nap1.

**Figure S3.**
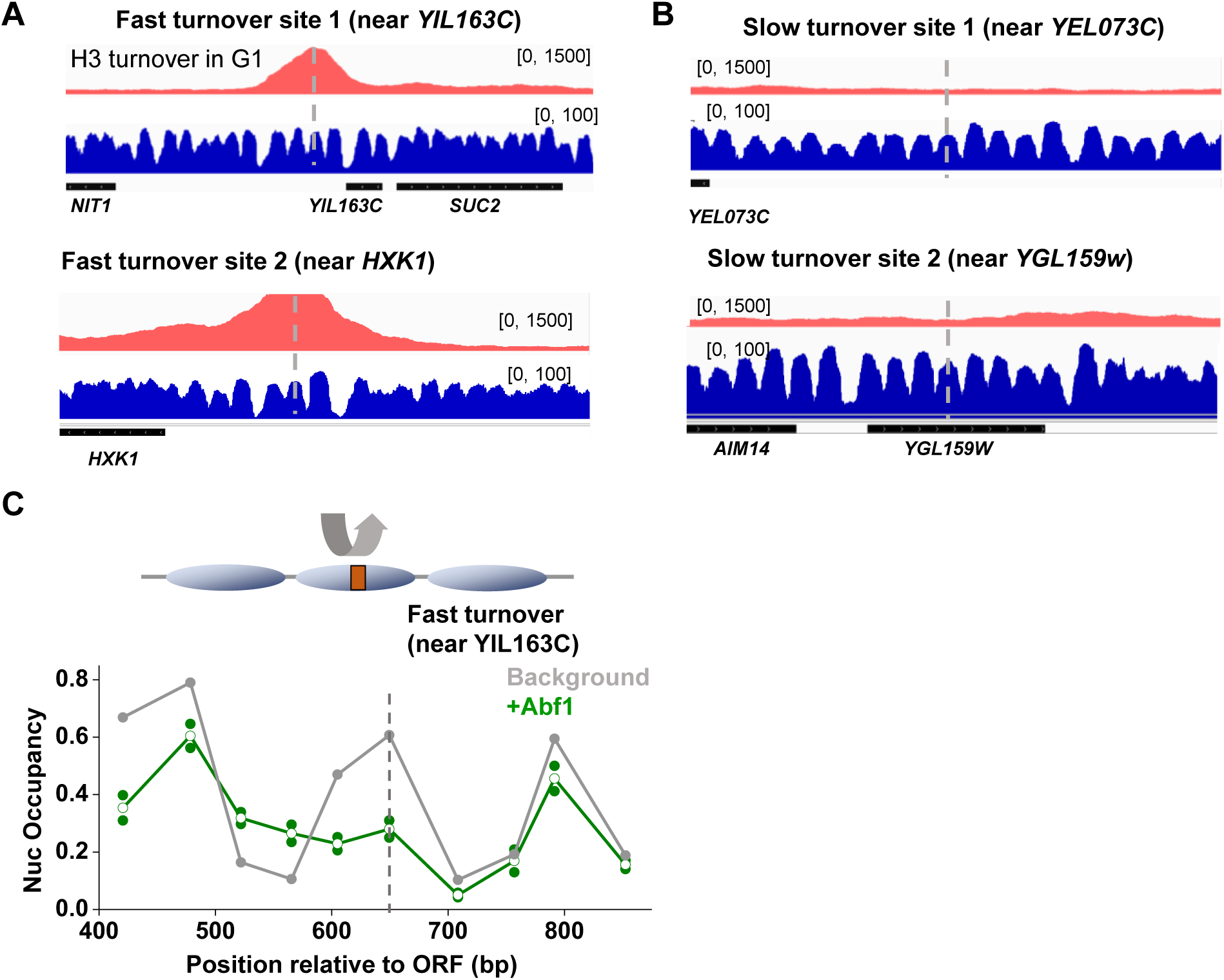
Histone turnover data and Abf1 invasion, related to Figure 4. **A)** IgV tracks showing histone turnover (red) and nucleosome occupancy (blue) near the two fast turnover sites used in Figure 4B. The histone turnover data are from Kassem et al., and the Mnase-seq data are generated by our own lab. The dashed lines mark the locations of LexO. **B)** Same as in A except for the two slow turnover sites used in Figure 4C. **C)** Nucleosome occupany near an Abf1 motif engineered into a fast turnover nucleosome (same site as the left panel in Figure 4B). Nucleosome in this case is displaced by the endogenous Abf1.

**Figure S4.**
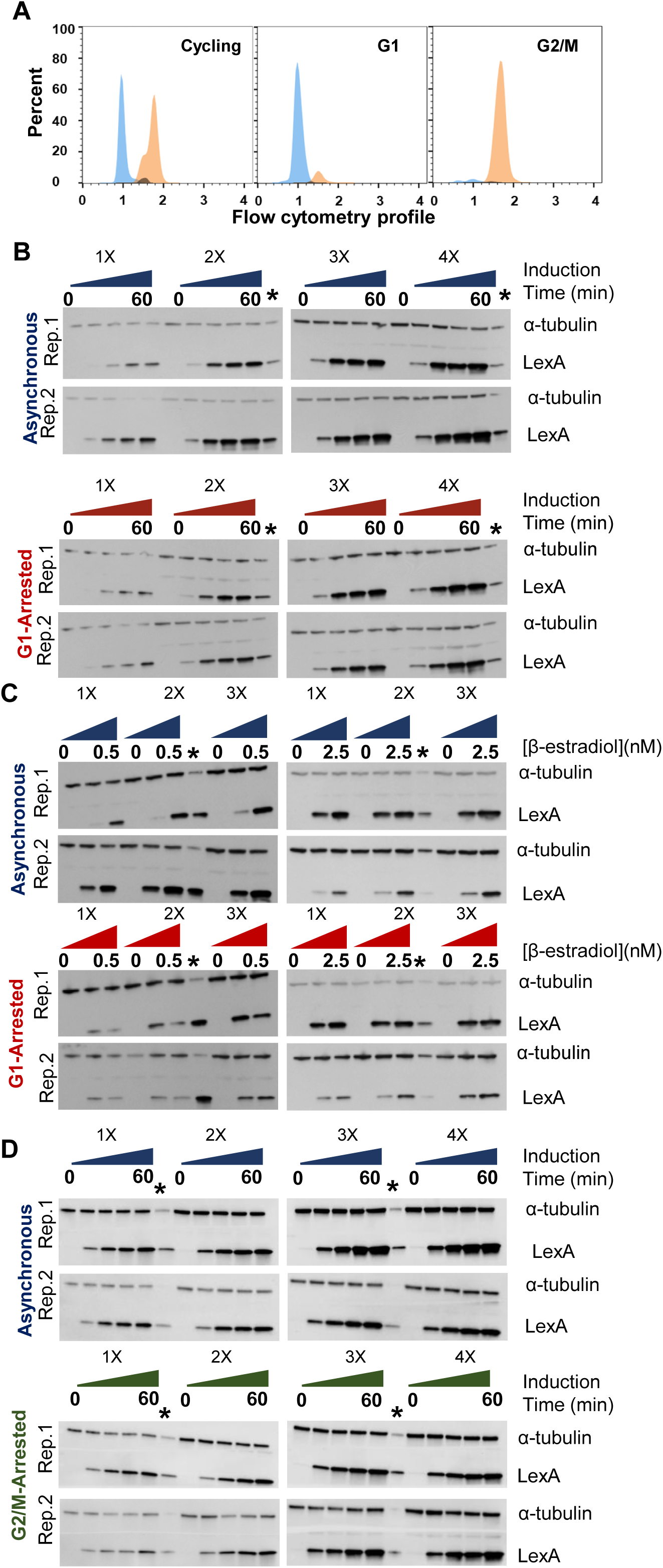
Cell cycle arrest and quantification of LexA induction levels, relate to Figure 5. **A)** Flow cytometric analysis of the DNA content in cycling cells (left), cells arrest in G1 by α factor (middle) and in G2/M by nocodazole (right). **B)** Western blot of LexA induction in cycling vs G1-arrested cells. **C)** Western blot of LexA induction in response to variable amount of β-estradiol in cycling vs G1-arrested cells. **D)** Western blot of LexA induction in cycling vs G2/M-arrested cells.

**Figure S5.**
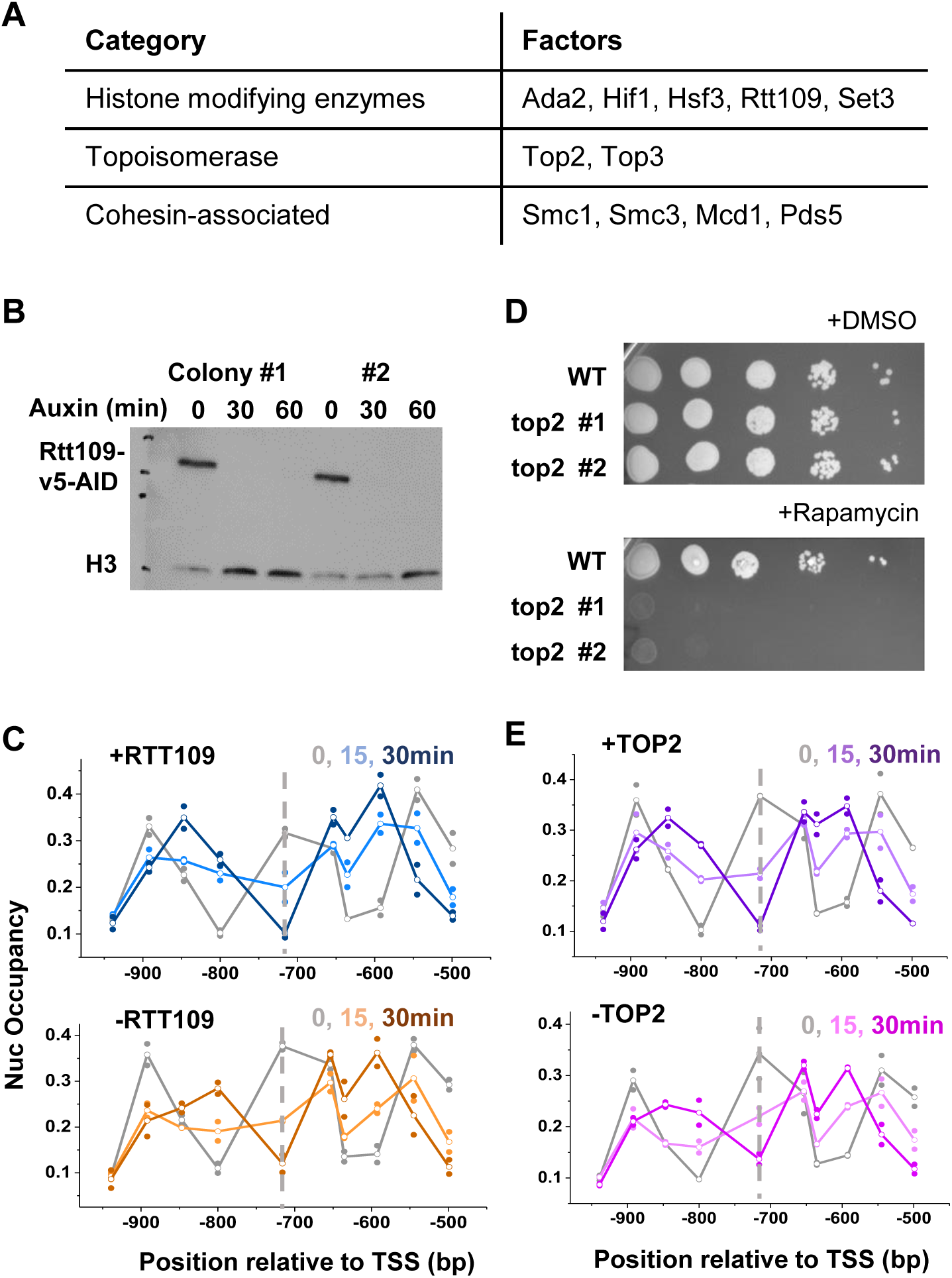
Nucleosome invasion by LexA is not affected by Top2 and Rtt109, related to Figure 6. **A)** List of chromatin-related factors that are transcriptionally repressed in G1. **B)** Western blot confirming auxin-induced degradation of Rtt109. **C)** Time course of LexA invasion into nucleosome –4 in cells with or without Rtt109. **D)** Spotting assay confirming the lethality of cells with Top2 anchor away. **E)** Time course of LexA invasion into nucleosome –4 in cells with or without Top2.

**Figure S6.**
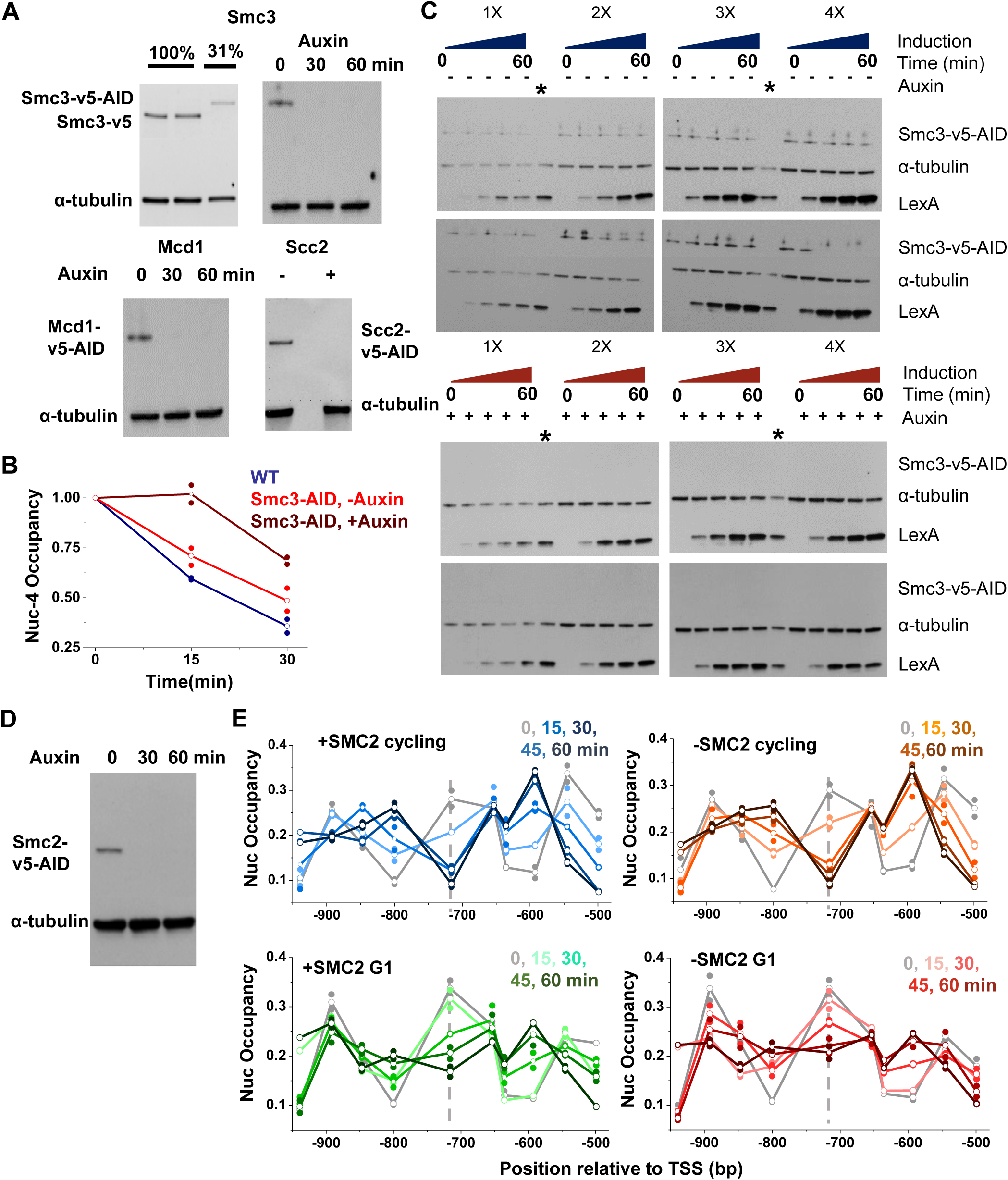
LexA expression and nucleosome invasion in the absence of cohesin or condensin, related to Figure 6. **A)** Western blots confirming auxin-induced degradation of Smc3, Mcd1, and Scc2. For Smc3, we also compared the WT level (Smc3-v5) to Smc3-v5-AID with no Auxin (upper left panel), which shows significant basal degradation. **B)** Nucleosome occupancy change over the LexO site upon LexA induction in WT and Smc3-AID ±auxin conditions. **C)** Western blot measuring LexA induction level in the presence and absence of Smc3. **D)** Western blot confirming auxin-induced degradation of Smc2. **D)** Time course of LexA invasion into nucleosome –4 in Smc2-AID strain ±auxin in cycling vs G1-arrested cells.

**Figure S7.**
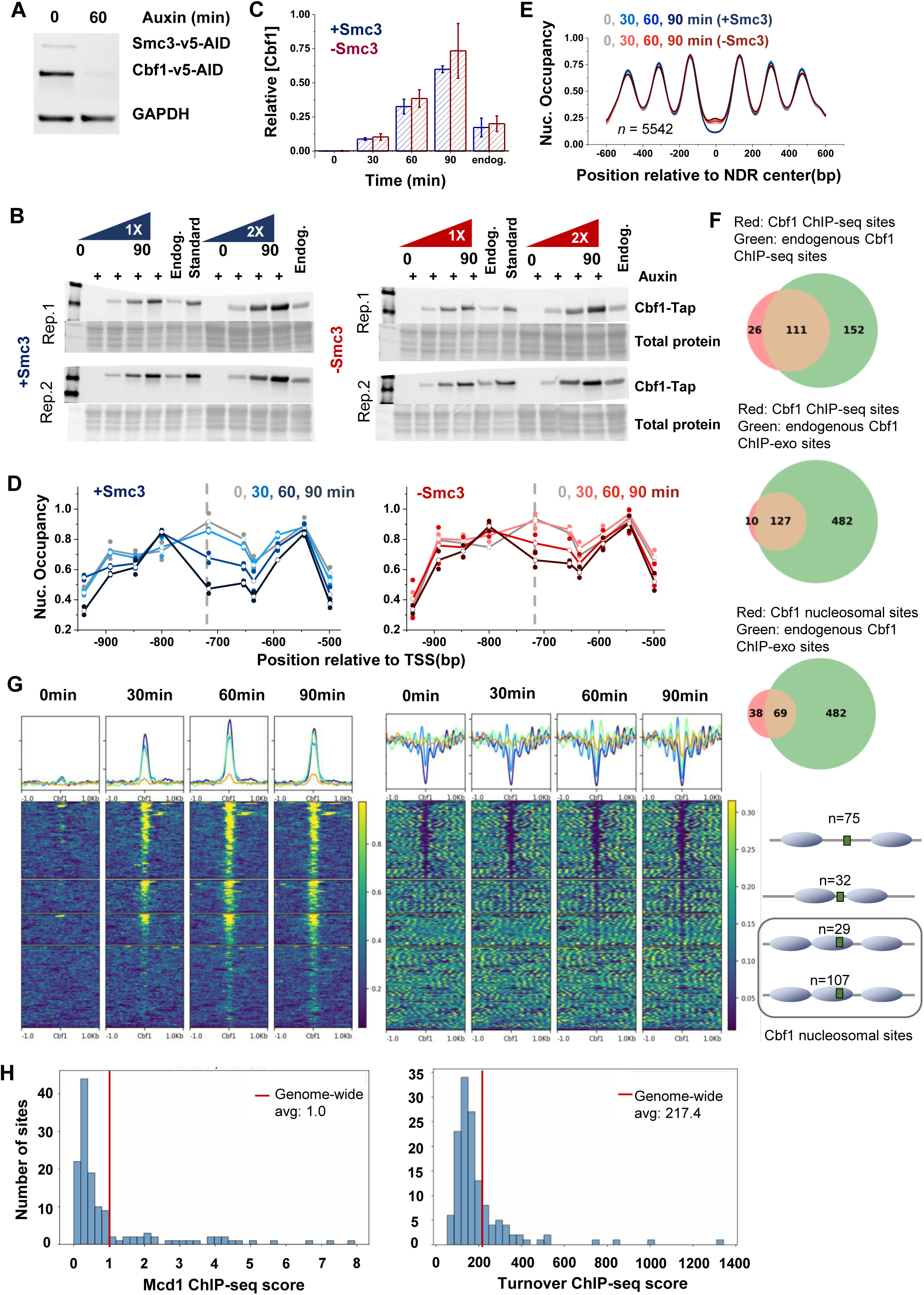
Cbf1 binding and nucleosome invasion in the presence or absence of Smc3, related to Figure 7. **A)** Western blot confirming auxin-induced degradation of Smc3 and Cbf1. **B)** Western blot analysis of Cbf1 induction levels in the presence or absence of Smc3. Total protein levels were used as loading controls (see Methods). **C)** Quantification of Cbf1 expression levels following induction in WT versus Smc3-depleted cells, and comparison with endogenous Cbf1 level. **D)** Time course of Cbf1 invasion into nucleosome –4 in the presence or absence of Smc3. **E)** Average nucleosome occupancy profile near genome-wide NDRs, aligned at NDR centers, in ± Smc3 cells at all time points of Cbf1 induction. **F)** Overlap of Cbf1 binding sites detected in this study with previously published datasets. **G)** ChIP-seq (left) and MNase-seq (right) over different classes of Cbf1 binding sites. From top to bottom: 1) sites in wide NDRs, 2) sites in narrow NDRs or nucleosome edges, 3) sites embedded within nucleosomes with strong Cbf1 ChIP-seq signals, and 4) sites within nucleosomes with weak Cbf1 ChIP-seq signals. The last two groups are combined as nucleosomal sites. **H)** Mcd1 ChIP-seq signals (left) and H3 turnover signals (right) at nucleosome-embedded Cbf1 sites. Genome-wide averages are marked by vertical bars and used as thresholds to group sites into high- versus low-signal categories.

## REFERENCES

1. Pina, B., Bruggemeier, U., and Beato, M. (1990). Nucleosome positioning modulates accessibility of regulatory proteins to the mouse mammary tumor virus promoter. Cell 60, 719–731. 10.1016/0092-8674(90)90087-u.

2. Polach, K.J., and Widom, J. (1995). Mechanism of protein access to specific DNA sequences in chromatin: a dynamic equilibrium model for gene regulation. J Mol Biol 254, 130–149. 10.1006/jmbi.1995.0606.

3. Luo, Y., North, J.A., Rose, S.D., and Poirier, M.G. (2014). Nucleosomes accelerate transcription factor dissociation. Nucleic Acids Res 42, 3017–3027. 10.1093/nar/gkt1319.

4. Cirillo, L.A., Lin, F.R., Cuesta, I., Friedman, D., Jarnik, M., and Zaret, K.S. (2002). Opening of compacted chromatin by early developmental transcription factors HNF3 (FoxA) and GATA-4. Mol Cell 9, 279–289. 10.1016/s1097-2765(02)00459-8.

5. Fernandez Garcia, M., Moore, C.D., Schulz, K.N., Alberto, O., Donague, G., Harrison, M.M., Zhu, H., and Zaret, K.S. (2019). Structural Features of Transcription Factors Associating with Nucleosome Binding. Mol Cell 75, 921–932 e926. 10.1016/j.molcel.2019.06.009.

6. Soufi, A., Garcia, M.F., Jaroszewicz, A., Osman, N., Pellegrini, M., and Zaret, K.S. (2015). Pioneer transcription factors target partial DNA motifs on nucleosomes to initiate reprogramming. Cell 161, 555–568. 10.1016/j.cell.2015.03.017.

7. Jung, J., Zheng, M., Goldfarb, M., and Zaret, K.S. (1999). Initiation of mammalian liver development from endoderm by fibroblast growth factors. Science 284, 1998–2003. 10.1126/science.284.5422.1998.

8. Tapia, N., MacCarthy, C., Esch, D., Gabriele Marthaler, A., Tiemann, U., Arauzo-Bravo, M.J., Jauch, R., Cojocaru, V., and Scholer, H.R. (2015). Dissecting the role of distinct OCT4-SOX2 heterodimer configurations in pluripotency. Sci Rep 5, 13533. 10.1038/srep13533.

9. Bulyk, M.L., Drouin, J., Harrison, M.M., Taipale, J., and Zaret, K.S. (2023). Pioneer factors - key regulators of chromatin and gene expression. Nat Rev Genet 24, 809–815. 10.1038/s41576-023-00648-z.

10. Stoeber, S., Godin, H., Xu, C., and Bai, L. (2024). Pioneer factors: nature or nurture? Crit Rev Biochem Mol Biol 59, 139–153. 10.1080/10409238.2024.2355885.

11. Donovan, B.T., Chen, H., Jipa, C., Bai, L., and Poirier, M.G. (2019). Dissociation rate compensation mechanism for budding yeast pioneer transcription factors. Elife 8. 10.7554/eLife.43008.

12. McDaniel, S.L., Gibson, T.J., Schulz, K.N., Fernandez Garcia, M., Nevil, M., Jain, S.U., Lewis, P.W., Zaret, K.S., and Harrison, M.M. (2019). Continued Activity of the Pioneer Factor Zelda Is Required to Drive Zygotic Genome Activation. Mol Cell 74, 185–195 e184. 10.1016/j.molcel.2019.01.014.

13. Donovan, B.T., Chen, H., Eek, P., Meng, Z., Jipa, C., Tan, S., Bai, L., and Poirier, M.G. (2023). Basic helix-loop-helix pioneer factors interact with the histone octamer to invade nucleosomes and generate nucleosome-depleted regions. Mol Cell 83, 1251–1263 e1256. 10.1016/j.molcel.2023.03.006.

14. Feng, X.A., Yamadi, M., Fu, Y., Ness, K.M., Liu, C., Ahmed, I., Bowman, G.D., Johnson, M.E., Ha, T., and Wu, C. (2025). GAGA zinc finger transcription factor searches chromatin by 1D-3D facilitated diffusion. Nat Struct Mol Biol. 10.1038/s41594-025-01643-0.

15. (!!! INVALID CITATION !!! 15-17).

16. Yan, C., Chen, H., and Bai, L. (2018). Systematic Study of Nucleosome-Displacing Factors in Budding Yeast. Mol Cell 71, 294–305 e294. 10.1016/j.molcel.2018.06.017.

17. Horisawa, K., Udono, M., Ueno, K., Ohkawa, Y., Nagasaki, M., Sekiya, S., and Suzuki, A. (2020). The Dynamics of Transcriptional Activation by Hepatic Reprogramming Factors. Mol Cell 79, 660–676 e668. 10.1016/j.molcel.2020.07.012.

18. Lerner, J., Gomez-Garcia, P.A., McCarthy, R.L., Liu, Z., Lakadamyali, M., and Zaret, K.S. (2020). Two-Parameter Mobility Assessments Discriminate Diverse Regulatory Factor Behaviors in Chromatin. Mol Cell 79, 677–688 e676. 10.1016/j.molcel.2020.05.036.

19. Hansen, J.L., Loell, K.J., and Cohen, B.A. (2022). A test of the pioneer factor hypothesis using ectopic liver gene activation. Elife 11. 10.7554/eLife.73358.

20. Hansen, J.L., and Cohen, B.A. (2022). A quantitative metric of pioneer activity reveals that HNF4A has stronger in vivo pioneer activity than FOXA1. Genome Biol 23, 221. 10.1186/s13059-022-02792-x.

21. Gaffney, D.J., McVicker, G., Pai, A.A., Fondufe-Mittendorf, Y.N., Lewellen, N., Michelini, K., Widom, J., Gilad, Y., and Pritchard, J.K. (2012). Controls of nucleosome positioning in the human genome. PLoS Genet 8, e1003036. 10.1371/journal.pgen.1003036.

22. Zhang, A.P., Pigli, Y.Z., and Rice, P.A. (2010). Structure of the LexA-DNA complex and implications for SOS box measurement. Nature 466, 883–886. 10.1038/nature09200.

23. Li, G., and Widom, J. (2004). Nucleosomes facilitate their own invasion. Nat Struct Mol Biol 11, 763–769. 10.1038/nsmb801.

24. Chen, R.W., Stoeber, S.D., Nodelman, I.M., Chen, H., Yang, W., Bowman, G.D., Bai, L., and Poirier, M.G. (2025). Native nucleosome-positioning elements as alternatives to the 601 sequence for nucleosome repositioning studies. Nucleic Acids Res 53. 10.1093/nar/gkaf822.

25. Chen, H., Kharerin, H., Dhasarathy, A., Kladde, M., and Bai, L. (2022). Partitioned usage of chromatin remodelers by nucleosome-displacing factors. Cell Rep 40, 111250. 10.1016/j.celrep.2022.111250.

26. Zhang, Q., Yoon, Y., Yu, Y., Parnell, E.J., Garay, J.A., Mwangi, M.M., Cross, F.R., Stillman, D.J., and Bai, L. (2013). Stochastic expression and epigenetic memory at the yeast HO promoter. Proc Natl Acad Sci U S A 110, 14012–14017. 10.1073/pnas.1306113110.

27. Zhou, K., Gaullier, G., and Luger, K. (2019). Nucleosome structure and dynamics are coming of age. Nat Struct Mol Biol 26, 3–13. 10.1038/s41594-018-0166-x.

28. Li, G., Levitus, M., Bustamante, C., and Widom, J. (2005). Rapid spontaneous accessibility of nucleosomal DNA. Nat Struct Mol Biol 12, 46–53. 10.1038/nsmb869.

29. Clapier, C.R., Iwasa, J., Cairns, B.R., and Peterson, C.L. (2017). Mechanisms of action and regulation of ATP-dependent chromatin-remodelling complexes. Nat Rev Mol Cell Biol 18, 407–422. 10.1038/nrm.2017.26.

30. Saha, A., Wittmeyer, J., and Cairns, B.R. (2002). Chromatin remodeling by RSC involves ATP-dependent DNA translocation. Genes Dev 16, 2120–2134. 10.1101/gad.995002.

31. Ganguli, D., Chereji, R.V., Iben, J.R., Cole, H.A., and Clark, D.J. (2014). RSC-dependent constructive and destructive interference between opposing arrays of phased nucleosomes in yeast. Genome Res 24, 1637–1649. 10.1101/gr.177014.114.

32. Kubik, S., Bruzzone, M.J., Challal, D., Dreos, R., Mattarocci, S., Bucher, P., Libri, D., and Shore, D. (2019). Opposing chromatin remodelers control transcription initiation frequency and start site selection. Nat Struct Mol Biol 26, 744–754. 10.1038/s41594-019-0273-3.

33. Brahma, S., and Henikoff, S. (2019). RSC-Associated Subnucleosomes Define MNase-Sensitive Promoters in Yeast. Mol Cell 73, 238–249 e233. 10.1016/j.molcel.2018.10.046.

34. Hammond, C.M., Stromme, C.B., Huang, H., Patel, D.J., and Groth, A. (2017). Histone chaperone networks shaping chromatin function. Nat Rev Mol Cell Biol 18, 141–158. 10.1038/nrm.2016.159.

35. Zhou, K., Liu, Y., and Luger, K. (2020). Histone chaperone FACT FAcilitates Chromatin Transcription: mechanistic and structural insights. Curr Opin Struct Biol 65, 26–32. 10.1016/j.sbi.2020.05.019.

36. Jeronimo, C., Poitras, C., and Robert, F. (2019). Histone Recycling by FACT and Spt6 during Transcription Prevents the Scrambling of Histone Modifications. Cell Rep 28, 1206–1218 e1208. 10.1016/j.celrep.2019.06.097.

37. Formosa, T., and Winston, F. (2020). The role of FACT in managing chromatin: disruption, assembly, or repair? Nucleic Acids Res 48, 11929–11941. 10.1093/nar/gkaa912.

38. McCullough, L., Rawlins, R., Olsen, A., Xin, H., Stillman, D.J., and Formosa, T. (2011). Insight into the mechanism of nucleosome reorganization from histone mutants that suppress defects in the FACT histone chaperone. Genetics 188, 835–846. 10.1534/genetics.111.128769.

39. Schlesinger, M.B., and Formosa, T. (2000). POB3 is required for both transcription and replication in the yeast Saccharomyces cerevisiae. Genetics 155, 1593–1606. 10.1093/genetics/155.4.1593.

40. Jamai, A., Imoberdorf, R.M., and Strubin, M. (2007). Continuous histone H2B and transcription-dependent histone H3 exchange in yeast cells outside of replication. Mol Cell 25, 345–355. 10.1016/j.molcel.2007.01.019.

41. Dion, M.F., Kaplan, T., Kim, M., Buratowski, S., Friedman, N., and Rando, O.J. (2007). Dynamics of replication-independent histone turnover in budding yeast. Science 315, 1405–1408. 10.1126/science.1134053.

42. Kassem, S., Ferrari, P., Hughes, A.L., Soudet, J., Rando, O.J., and Strubin, M. (2020). Histone exchange is associated with activator function at transcribed promoters and with repression at histone loci. Sci Adv 6. 10.1126/sciadv.abb0333.

43. Neely, K.E., Hassan, A.H., Wallberg, A.E., Steger, D.J., Cairns, B.R., Wright, A.P., and Workman, J.L. (1999). Activation domain-mediated targeting of the SWI/SNF complex to promoters stimulates transcription from nucleosome arrays. Mol Cell 4, 649–655. 10.1016/s1097-2765(00)80216-6.

44. Han, J., Zhou, H., Horazdovsky, B., Zhang, K., Xu, R.M., and Zhang, Z. (2007). Rtt109 acetylates histone H3 lysine 56 and functions in DNA replication. Science 315, 653–655. 10.1126/science.1133234.

45. Neumann, H., Hancock, S.M., Buning, R., Routh, A., Chapman, L., Somers, J., Owen-Hughes, T., van Noort, J., Rhodes, D., and Chin, J.W. (2009). A method for genetically installing site-specific acetylation in recombinant histones defines the effects of H3 K56 acetylation. Mol Cell 36, 153–163. 10.1016/j.molcel.2009.07.027.

46. Teves, S.S., and Henikoff, S. (2014). Transcription-generated torsional stress destabilizes nucleosomes. Nat Struct Mol Biol 21, 88–94. 10.1038/nsmb.2723.

47. Liu, L.F., and Wang, J.C. (1987). Supercoiling of the DNA template during transcription. Proc Natl Acad Sci U S A 84, 7024–7027. 10.1073/pnas.84.20.7024.

48. Yan, J., Enge, M., Whitington, T., Dave, K., Liu, J., Sur, I., Schmierer, B., Jolma, A., Kivioja, T., Taipale, M., and Taipale, J. (2013). Transcription factor binding in human cells occurs in dense clusters formed around cohesin anchor sites. Cell 154, 801–813. 10.1016/j.cell.2013.07.034.

49. Spellman, P.T., Sherlock, G., Zhang, M.Q., Iyer, V.R., Anders, K., Eisen, M.B., Brown, P.O., Botstein, D., and Futcher, B. (1998). Comprehensive identification of cell cycle-regulated genes of the yeast Saccharomyces cerevisiae by microarray hybridization. Mol Biol Cell 9, 3273–3297. 10.1091/mbc.9.12.3273.

50. Uhlmann, F., Lottspeich, F., and Nasmyth, K. (1999). Sister-chromatid separation at anaphase onset is promoted by cleavage of the cohesin subunit Scc1. Nature 400, 37–42. 10.1038/21831.

51. Schalbetter, S.A., Goloborodko, A., Fudenberg, G., Belton, J.M., Miles, C., Yu, M., Dekker, J., Mirny, L., and Baxter, J. (2017). SMC complexes differentially compact mitotic chromosomes according to genomic context. Nat Cell Biol 19, 1071–1080. 10.1038/ncb3594.

52. Zou, F., Li, Y., Foldes, T., Pinholt, H.D., Smith, C., Mirny, L., and Bai, L. (2025). Condensin Accelerates Long-Range Intra-Chromosomal Interactions. bioRxiv. 10.1101/2025.05.02.651983.

53. Lee, J., Simpson, L., Li, Y., Becker, S., Zou, F., Zhang, X., and Bai, L. (2024). Transcription Factor Condensates Mediate Clustering of MET Regulon and Enhancement in Gene Expression. bioRxiv. 10.1101/2024.02.06.579062.

54. Rossi, M.J., Kuntala, P.K., Lai, W.K.M., Yamada, N., Badjatia, N., Mittal, C., Kuzu, G., Bocklund, K., Farrell, N.P., Blanda, T.R., et al. (2021). A high-resolution protein architecture of the budding yeast genome. Nature 592, 309–314. 10.1038/s41586-021-03314-8.

55. Costantino, L., Hsieh, T.S., Lamothe, R., Darzacq, X., and Koshland, D. (2020). Cohesin residency determines chromatin loop patterns. Elife 9. 10.7554/eLife.59889.

56. Tang, X., Li, T., Liu, S., Wisniewski, J., Zheng, Q., Rong, Y., Lavis, L.D., and Wu, C. (2022). Kinetic principles underlying pioneer function of GAGA transcription factor in live cells. Nat Struct Mol Biol 29, 665–676. 10.1038/s41594-022-00800-z.

57. Eustermann, S., Schall, K., Kostrewa, D., Lakomek, K., Strauss, M., Moldt, M., and Hopfner, K.P. (2018). Structural basis for ATP-dependent chromatin remodelling by the INO80 complex. Nature 556, 386–390. 10.1038/s41586-018-0029-y.

58. Wagner, F.R., Dienemann, C., Wang, H., Stutzer, A., Tegunov, D., Urlaub, H., and Cramer, P. (2020). Structure of SWI/SNF chromatin remodeller RSC bound to a nucleosome. Nature 579, 448–451. 10.1038/s41586-020-2088-0.

59. Kubik, S., Bruzzone, M.J., Jacquet, P., Falcone, J.L., Rougemont, J., and Shore, D. (2015). Nucleosome Stability Distinguishes Two Different Promoter Types at All Protein-Coding Genes in Yeast. Mol Cell 60, 422–434. 10.1016/j.molcel.2015.10.002.

60. Kraushaar, D.C., Jin, W., Maunakea, A., Abraham, B., Ha, M., and Zhao, K. (2013). Genome-wide incorporation dynamics reveal distinct categories of turnover for the histone variant H3.3. Genome Biol 14, R121. 10.1186/gb-2013-14-10-r121.

61. Roberts, G.A., Ozkan, B., Gachulincova, I., O’Dwyer, M.R., Hall-Ponsele, E., Saxena, M., Robinson, P.J., and Soufi, A. (2021). Dissecting OCT4 defines the role of nucleosome binding in pluripotency. Nat Cell Biol 23, 834–845. 10.1038/s41556-021-00727-5.

62. Lerner, J., Katznelson, A., Zhang, J., and Zaret, K.S. (2023). Different chromatin-scanning modes lead to targeting of compacted chromatin by pioneer factors FOXA1 and SOX2. Cell Rep 42, 112748. 10.1016/j.celrep.2023.112748.

63. Gibson, T.J., Larson, E.D., and Harrison, M.M. (2024). Protein-intrinsic properties and context-dependent effects regulate pioneer factor binding and function. Nat Struct Mol Biol 31, 548–558. 10.1038/s41594-024-01231-8.

64. Noutsou, M., Li, J., Ling, J., Jones, J., Wang, Y., Chen, Y., and Sen, G.L. (2017). The Cohesin Complex Is Necessary for Epidermal Progenitor Cell Function through Maintenance of Self-Renewal Genes. Cell Rep 20, 3005–3013. 10.1016/j.celrep.2017.09.003.

65. Yamin, K., Bijlani, S., Berman, J., Soni, A., Shlomai, J., Buragohain, B.M., Werbner, M., Gal-Tanamy, M., Matityahu, A., and Onn, I. (2022). Fold-change of chromatin condensation in yeast is a conserved property. Sci Rep 12, 17393. 10.1038/s41598-022-22340-8.

66. Bird, A. (2025). Cohesin as an essential disruptor of chromosome organization. Mol Cell 85, 1054–1057. 10.1016/j.molcel.2025.01.010.

67. Davidson, I.F., Barth, R., Nagasaka, K., Tang, W., Wutz, G., Horn, S., Janissen, R., Stocsits, R.R., Chlosta, E., Bauer, B.W., et al. (2025). Cohesin supercoils DNA during loop extrusion. Cell Rep 44, 115856. 10.1016/j.celrep.2025.115856.

68. Du, M., Zou, F., Li, Y., Yan, Y., and Bai, L. (2022). Chemically Induced Chromosomal Interaction (CICI) method to study chromosome dynamics and its biological roles. Nat Commun 13, 757. 10.1038/s41467-022-28416-3.

69. Rosebrock, A.P. (2017). Analysis of the Budding Yeast Cell Cycle by Flow Cytometry. Cold Spring Harb Protoc 2017. 10.1101/pdb.prot088740.

70. Bai, L., Ondracka, A., and Cross, F.R. (2011). Multiple sequence-specific factors generate the nucleosome-depleted region on CLN2 promoter. Mol Cell 42, 465–476. 10.1016/j.molcel.2011.03.028.

71. Jorgensen, P., Edgington, N.P., Schneider, B.L., Rupes, I., Tyers, M., and Futcher, B. (2007). The size of the nucleus increases as yeast cells grow. Mol Biol Cell 18, 3523–3532. 10.1091/mbc.e06-10-0973.

72. Carrillo, E., Ben-Ari, G., Wildenhain, J., Tyers, M., Grammentz, D., and Lee, T.A. (2012). Characterizing the roles of Met31 and Met32 in coordinating Met4-activated transcription in the absence of Met30. Mol Biol Cell 23, 1928–1942. 10.1091/mbc.E11-06-0532.

73. Chen, K., Xi, Y., Pan, X., Li, Z., Kaestner, K., Tyler, J., Dent, S., He, X., and Li, W. (2013). DANPOS: dynamic analysis of nucleosome position and occupancy by sequencing. Genome Res 23, 341–351. 10.1101/gr.142067.112.

74. Henikoff, J.G., Belsky, J.A., Krassovsky, K., MacAlpine, D.M., and Henikoff, S. (2011). Epigenome characterization at single base-pair resolution. Proc Natl Acad Sci U S A 108, 18318–18323. 10.1073/pnas.1110731108.

75. Spivak, A.T., and Stormo, G.D. (2012). ScerTF: a comprehensive database of benchmarked position weight matrices for Saccharomyces species. Nucleic Acids Res 40, D162–168. 10.1093/nar/gkr1180.

